# Stable tug-of-war between kinesin-1 and cytoplasmic dynein upon different ATP and roadblock concentrations

**DOI:** 10.1101/2020.06.08.140806

**Authors:** Gina A. Monzon, Lara Scharrel, Ashwin DSouza, Ludger Santen, Stefan Diez

## Abstract

The maintenance of intracellular processes like organelle transport and cell division depend on bidirectional movement along microtubules. These processes typically require kinesin and dynein motor proteins which move with opposite directionality. Because both types of motors are often simultaneously bound to the cargo, regulatory mechanisms are required to ensure controlled directional transport. Recently, it has been shown that parameters like mechanical motor activation, ATP concentration and roadblocks on the microtubule surface differentially influence the activity of kinesin and dynein motors in distinct manners. However, how these parameters affect bidirectional transport systems has not been studied. Here, we investigate the regulatory influence of these three parameter using in vitro gliding motility assays and stochastic simulations. We find that the number of active kinesin and dynein motors determines the transport direction and velocity, but that variations in ATP concentration and roadblock density have no significant effect. Thus, factors influencing the force balance between opposite motors appear to be important, whereas the detailed stepping kinetics and bypassing capabilities of the motors have only little effect.

## INTRODUCTION

Intracellular transport is essential for the functionality of a cell. For instance, intracellular transport is used for chromosome segregation during cell division (Lodish et al., 2000; Verhey and Hammond, 2009) and dysfunction of intracellular transport leads to neurodegenerative diseases like Alzheimer or ALS (De Vos et al., 2008; Goldstein et al., 2001; Hurd et al., 1996; Chen et al., 2014; Karki et al., 1999). Intracellular transport needs an intracellular “street-network” which is provided by the microtubule cytoskeleton. On microtubules, cargo like mitochondria can be transported over long distances (Hollenbeck et al., 2005). Microtubule-based transport is carried out by teams of the opposite-directed motor proteins kinesin and dynein. These motors actively move cargoes backwards and forwards along microtubules, allowing them to be delivered to where they are needed. Mitochondria, for instance, are transported to locations of low ATP concentration (Morris et al., 1993) and chromosomes are perfectly aligned on spindle microtubules during cell division (She and Yang, 2017; Goshima et al., 2003). Thus, kinesin and dynein motors perform targeted intracellular transport. Importantly, however, teams of kinesin and dynein motors are known to be simultaneously bound to the cargo (Welte, 2004; Soppina et al., 2009; Hendricks et al., 2010). Consequently, having both opposite-directed motors attached to the cargo, it is unclear how targeted transport is ensured. Basically, the cargo could be transported in either the kinesin or the dynein direction. Without any regulatory mechanism, the cargo might even randomly change its direction or get stuck at a random position. However, the regulatory mechanisms ensuring targeted transport remain poorly understood.

In the past, different regulatory mechanisms have been proposed. One proposed mechanism suggests coordinating the motor activity to regulate bidirectional transport (Gross, 2004). In this model, motors are assumed to be a priori in a passive state. By activating one motor team, targeted cargo transport occurs in the direction of the active team (Gross, 2004). One such activation mechanism involves adaptor proteins (McKenney et. al., 2014; Schroeder et al., 2016; Blasius et al., 2007; Elshenawy et al., 2019). The adaptor protein dynactin plus a cargo adaptor, for instance, activates passive cytoplasmic dynein (McKenney et. al., 2014). Besides activation by adaptor proteins, dynein can also be mechanically activated (Monzon and Scharrel et al., 2019; Torisawa et al., 2014). Ally et al. (2009) have shown that a mechanical motor coupling is needed for cargo transport and De Rossi et al. (2017) hypothesize that motors might activate each other by exerting forces on the opposite motor team. Mechanical dynein activation has been shown to determine the velocity in unidirectional dynein-driven transport (Monzon and Scharrel et al., 2019). However, in bidirectional transport the role of mechanical dynein activation has not been considered in the past. For unidirectional dynein-driven transport Monzon and Scharrel et al. (2019) showed that mechanical dynein activation strongly depends on the number of involved dynein motors. This suggests that in bidirectional transport mechanical activation might also be linked to the number of motors. The influence of varying the number of motors has been studied before. Rezaul et al. (2016), for instance, reversed a dynein-driven membrane organelle in vivo by adding a large number of kinesin motors. Moreover, Vale et al. (1992) showed that the transport direction in bidirectional gliding assays depends on the number of kinesin motors. However, for a complete understanding of the role of mechanical activation a systematic analysis is needed.

Another model proposes modifying motor properties as a regulation mechanism for bidirectional transport (Müller et al., 2008). Modification of motor properties has been shown to lead to different motility states (Müller et al., 2008). One important motor property is the motor velocity. The single motor velocity can be modified by changing the ATP concentration (Schnitzer et al., 2000; Ross et al, 2006; Torisawa et al., 2014). Imagine a cargo being transported in both directions with identical velocities and frequent directional changes. If then an increase of the ATP concentration asymmetrically modifies the velocity of the motor teams, the team with the higher velocity might take over. Another motor property influenced by ATP concentration is the motor stall force (Mallik et al., 2004; Visscher et al., 1999). While dynein stall force increases linearly with ATP concentration (Mallik et al., 2004), kinesin stall force is slightly reduced for very low ATP concentrations but invariant for high ATP concentrations (Visscher et al., 1999). Depending on the stall force, Müller et al. (2008) defines the motor strength as the “stall force to detachment force ratio”. Motors having a large “stall force to detachment force ratio” are referred to as, strong motors” and motors with a small “stall force to detachment force ratio” as "weak motors” (Müller et al., 2008). Therefore, a higher dynein stall force would strengthen the dynein team. A higher dynein stall force could be reached by increasing the ATP concentration. As a consequence, with increasing ATP concentration, the dynein team might be able to reverse the transport direction of a kinesin-driven cargo. Consistent with this, a directional change as a function of the ATP concentration is predicted by the theoretical work of (Klein et al., 2014). However, changing the transport direction by the ATP concentration has not been tested experimentally.

Using hindering roadblocks or obstacles to regulate bidirectional transport is a third possible mechanism. Different reactions of single motor proteins have been observed when being hindered by a roadblock (Telley et al., 2009; Dixit et al., 2008; Schneider et al., 2015). While single kinesin detaches, for instance, when encountering the microtubule-associated protein tau, dynein continues stepping (Vershinin et al., 2007; Siahaan et al., 2019; Tan et al., 2019). Moreover, different mechanisms to bypass roadblocks have been observed in single-molecule experiments (Ferro et al., 2019). While kinesin has to detach and reattach behind the roadblock, dynein uses its ability to take sidesteps (Ferro et al., 2019). However, how a cargo that is bidirectionally transported by many motors reacts when encountering a roadblock remains unclear. The cargo might stop in front of the roadblock, detach, reverse its direction or continue walking. Moreover, the reaction might be different if the cargo was driven by dynein or kinesin before encountering a roadblock. Thus, whether roadblocks change the transport direction of bidirectionally transported cargo is unclear. Furthermore, the overall impact on the regulation of bidirectional transport would depend on the relative contribution of not only hindering roadblocks but also of other factors such as the number of motors and the ATP concentration.

To fully understand the relative contribution of different parameters to bidirectional transport a systematic approach is needed. Parameters such as number of motors needs to be varied systematically and the bidirectional transport needs to be measured without affecting the activity of the motors themselves. The use of microtubule gliding assays is one way to achieve this. In microtubule gliding assays, a coverslip (from now on referred to as surface) is coated with motors and a microtubule is freely placed above the surface. Motors in the region underneath the microtubule attach to the microtubule and step along it. Thereby the motors move the microtubule back and forth. The number of motors involved in the transport can be systematically changed by varying the motor density on the surface. To measure the collective behavior of motors without affecting the motors themselves, microtubules are labeled and tracked. Consequently, microtubule gliding assays (see Fig. 1a for an illustration) are highly suitable for systematically investigating the influence of the motor number on bidirectional transport. Moreover, ATP concentration in the surrounding can be systematically changed and microtubules can be coated with roadblocks at different concentrations. In this article, we perform a systematic analysis of the effects of motor number, ATP concentration and hindering roadblocks on bidirectional transport. Additionally, to get a detailed picture of the role of factors such as mechanical activation, we perform simulations of mathematical kinesin and dynein models. The models are based on all known single molecule parameters.

**Fig. 1.**
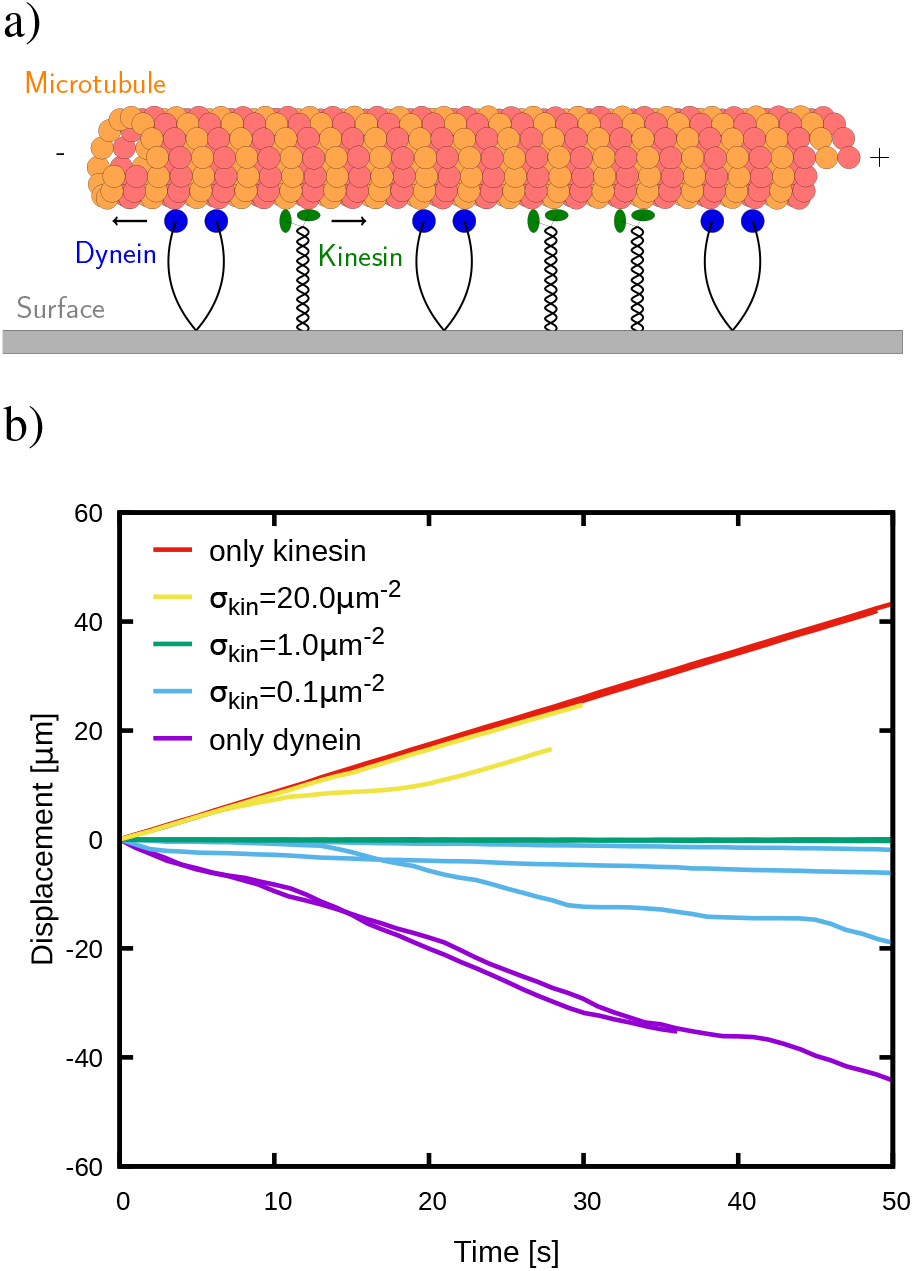
Varying the kinesin density regulates the transport direction in bidirectional microtubule gliding assays. a) Schematic diagram of a bidirectional gliding assay. Kinesin-1 and cytoplasmic dynein (from know on kinesin and dynein) are permanently bound to the surface (coverslip) with their tail region and attach to a microtubule situated above them. The density of motors determines the number of motors involved in the transport for a specific microtubule length. Attached motors move the microtubule back and forth when stepping on it. While attached kinesin steps towards the microtuble plus end, dynein steps towards the microtubule minus end. This results in a tug-of-war between the opposite directed motor teams. b) Example trajectories of microtubule gliding for varying kinesin densities and a constant dynein density of 64 μm^−2^. The microtubules of the shown trajectories have lengths within the interval (5 μm – 10 μm). Kinesin-driven transport is defined to be in positive direction (positive displacement) and dynein-driven transport in negative direction (negative displacement). For the “only kinesin” case (red curve) and a high kinesin density of 20 μm^−2^ (yellow curve) we see kinesin-driven transport. For a kinesin density of 1 μm^−2^ we see almost no net movement. Forces between the kinesin and the dynein team are balanced at this kinesin density. And for a very low kinesin density of 0.1 μm^−2^ (light blue curve) and the “only dynein” case (purple curve) we observe dynein-driven transport. Together we see three motility states: the kinesin-driven state, the balanced state and the dynein-driven state.

## RESULTS

### Regulating transport direction by varying the motor number

To determine the influence of motor number on bidirectional transport, we first systematically varied the motor density on the surface of the microtubule gliding assay. The motor density determines how many motors are underneath the microtubule and therefore determines the motor number involved in the transport process^1^. (See Fig. 1a for an illustration of the microtubule gliding assay). Previous studies showed that changing the kinesin number alters the transport direction (Rezaul et al., 2016; Vale et al., 1992). To test whether our gliding assay set-up also shows a directional change in dependence of the kinesin number, we first varied the kinesin density at a constant dynein density. Therefore, we coated the surface with conventional kinesin-1 and cytoplasmic mammalian dynein (hereafter referred to as kinesin and dynein). The kinesin density on the surface was varied from 0 to 100 μm^−2^ and the unidirectional cases "only dynein” and "only kinesin” were included as references. As a constant dynein density, a rather high dynein density of 64 μm^−2^ was applied to ensure directed dynein-transport^2^. To determine the transport direction, the microtubule plus end was labelled and gliding trajectories were measured. Example trajectories (Fig. 1b) show dynein-driven transport (purple and blue curves) for very low kinesin densities of 0.1 μm^−2^ and the "only dynein” case. For higher kinesin densities of 1 μm^−2^ almost no net movement was observed. In this case forces between the kinesin and dynein team are balanced and the transport is stuck. For higher kinesin densities of 20 μm^−2^ and in the "only kinesin” case kinesin-driven transport was observed. In total, three motility states could be distinguished depending on the kinesin density: (i) dynein-driven state, (ii) balanced state (almost no net movement) and (iii) kinesin-driven state. Thus, varying the kinesin density the transport direction can be changed.

When seeking to understand the regulation by the kinesin density and the role of mechanical dynein activation, an insight into the state of the motors at the molecular level is required. The number of attached motors and their internal state (active or passive) are important variables. Experimentally, it is difficult to determine the number of attached kinesin and dynein motors, and it is not possible to know whether the individual motors are active or passive. We therefore used mathematical models of single kinesin and dynein molecules developed by Monzon and Scharrel et al. (2019) and Klein et al. (2014). The models are based on all known kinesin and dynein single molecule properties. In Monzon and Scharrel et al. (2019), the models are used for unidirectional kinesin and dynein gliding assay simulations. Here, we adjusted the models to the bidirectional gliding assay set-up. An illustration of the bidirectional gliding assay simulation is shown in Fig. 2a and detailed model descriptions can be found in the Material and Methods section. Here, the bidirectional gliding assay simulations are used to obtain deeper insight to the molecular level.

**Fig. 2.**
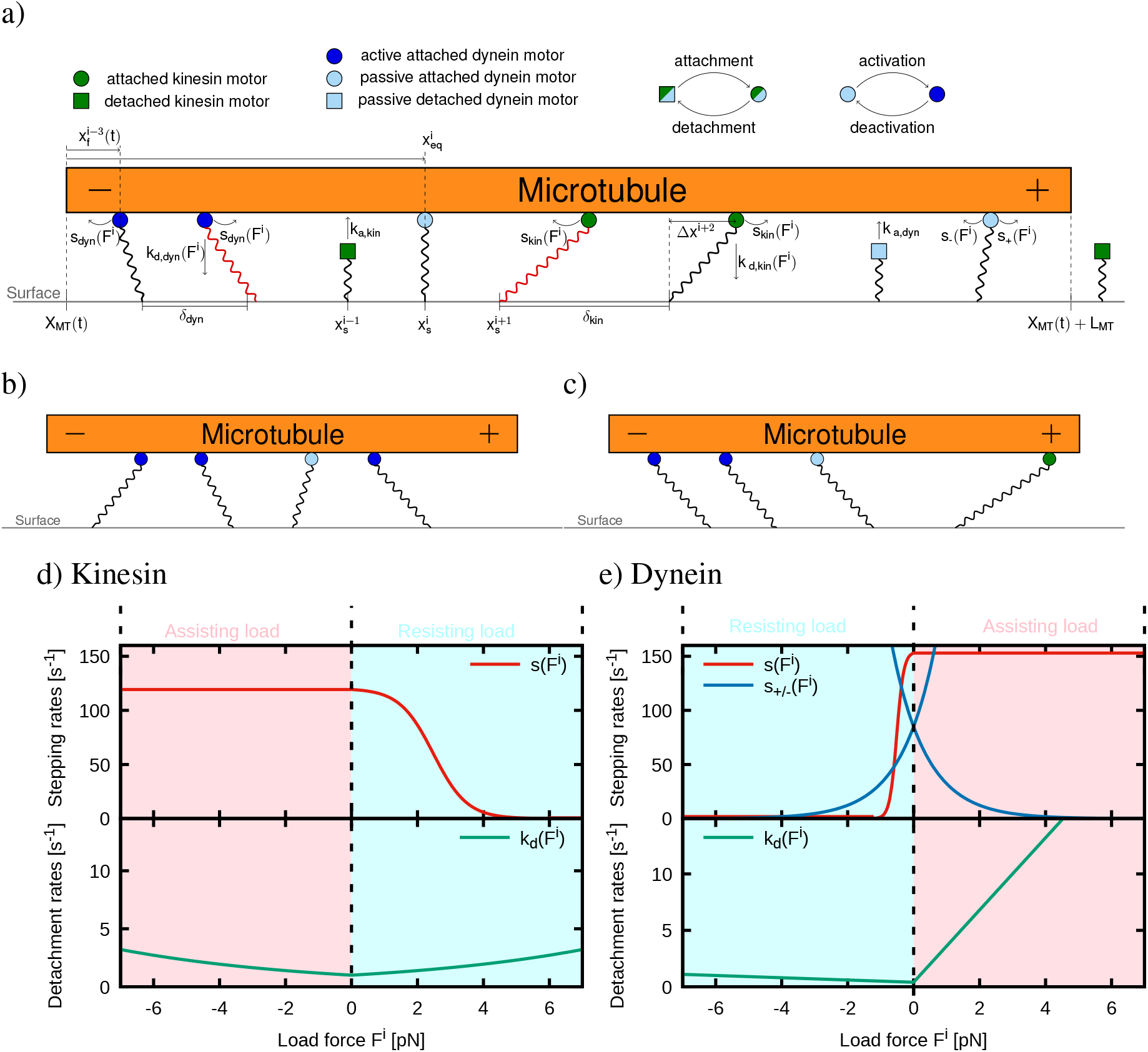
Using mathematical kinesin and dynein models for bidirectional microtubule gliding assays simulations give an insight to the molecular level of transport. a) Scheme of the bidirectional gliding assay simulation. The microtubule gliding assay is implemented as a one-dimensional system. In the one-dimensional simulation the microtubule position *X*_MT_(*t*) is determined by the microtubule minus end. The microtubule plus end is therefore at *X*_MT_(*t*) + *L*_MT_. with *L*_MT_ being the constant microtubule length. On the one-dimensional surface (coverslip), under the microtubule, kinesin and dynein motors are randomly distributed with mean distances *δ*_kin_ and *δ*_dyn_, respectively. The permanent position of the *i*^th^ motor on the surface is 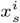 and, when attached, its position on the microtubule is 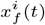 (filament position). Detached kinesin motors are drawn as green squares and attached kinesin motors as green circles. Detached dynein motors are drawn as light blue squares. Passive attached dynein is presented by light blue circles and active attached dynein by dark blue circles. While motors attach with the constant rates *k*_a,kin_ and *k*_a,dyn_, the detachment (*k*_d,kin_(*F^i^*) for kinesin and *k*_d,dyn_(*F^i^*) for passive and active dynein) and stepping (*s*_kin_(*F^i^*) for kinsein, *s*_dyn_(*F^i^*) for active dynein and *s*_±_(*F^i^*) for passive dynein) of attached motors depend on the load force of the motors. Since motors are modeled as linear springs, the load force of the motors is proportional to the motor deflection 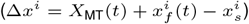. For load forces smaller than the stall force, kinesin and active dynein step pointedly towards the microtubule plus and minus end, respectively. For load forces greater than the motors stall force, kinesin and active dynein step backward. Passive dynein diffuses in the harmonic potential of the motors springs. b) Example motor deflections for the “only dynein” case. c) Alignment of attached dynein motors by the activity of one kinesin motor. d) Force dependent stepping (upper panel) and detachment rates (lower panel) for kinesin. Under forward load (assisting forces, negative for kinesin) the kinesin stepping (red curve) is high, but constant and decreases for backward loads (resisting forces, positive for kinesin) until reaching stall (*F*_s,kin_ = 6 pN). For a higher backward load than the stall force, kinesin steps backward with a small but constant rate. The detachment of kinesin (green curve, lower panel) increases exponentially and symmetrically for forward and backward load. e) Force dependent stepping (upper panel) and detachment rates (lower panel) for dynein. Active dynein steps with an high but constant rate (red curve) under forward load (assisting forces, positive for dynein). Under backward load (resisting forces, negative for dynein) the directional stepping of active dynein decreases until reaching the stall force (*F*_s,dyn_ = 1.25 pN). Beyond the stall force, active dynein steps backwards with a small but constant rate. The diffusive stepping of passive dynein (blue curve) increases exponentially for stepping towards the equilibrium position 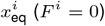 of the motor and decreases exponentially for stepping away from the equilibrium position. Thus *s*_±_(*F^i^*) is mirrored at the y-axis. The dynein detachment (green curve) increases linearly but asymmetrically for backward and forward load. Under forward load the detachment rate increases faster than under backward load. For more details regarding the model see Material and Methods chapter and (Monzon and Scharrel et. al. 2019).

To be able to rely on the information provided by the bidirectional gliding assay simulation, it has to be shown that the simulation reproduces the experimental observations. Therefore, velocity histograms from simulation and experiment were compared (see Fig. 3). For the histograms again, the kinesin densities were varied at a constant high dynein density. As defined before positive velocities represent kinesin-driven transport and negative velocities dynein-driven transport. In both, simulation and experiment a small peak was measured at high positive velocities for the “only kinesin” case and a high kinesin density (100 μm^−2^) (see Fig. 3). A widening of the peak towards lower positive velocities could be seen for intermediate kinesin density (20 μm^−2^). For low kinesin densities (1 μm^−2^), a peak was observed around zero, which is a bit wider for the simulation than for the experiment. For both the simulation and the experiment the peak then widened towards negative velocities for even lower kinesin densities (0.1 μm^−2^). For the “only dynein” case, simulation and experiment showed a broad distribution of negative velocities. Altogether for both simulation and experiment kinesin-driven transport was observed for high and intermediate kinesin densities, a balanced state at low kinesin density and dynein-driven transport for very low kinesin densities. Thus, the simulation reproduces the experiment reasonably well and can be considered a reliable tool to obtain a deeper insight to the bidirectional transport on a molecular level. Taken together these findings suggest that bidirectional transport strongly depends on the kinesin density. As a consequence, the number of kinesin motors can be used to regulate the directionality of bidirectional transport.

**Fig. 3.**
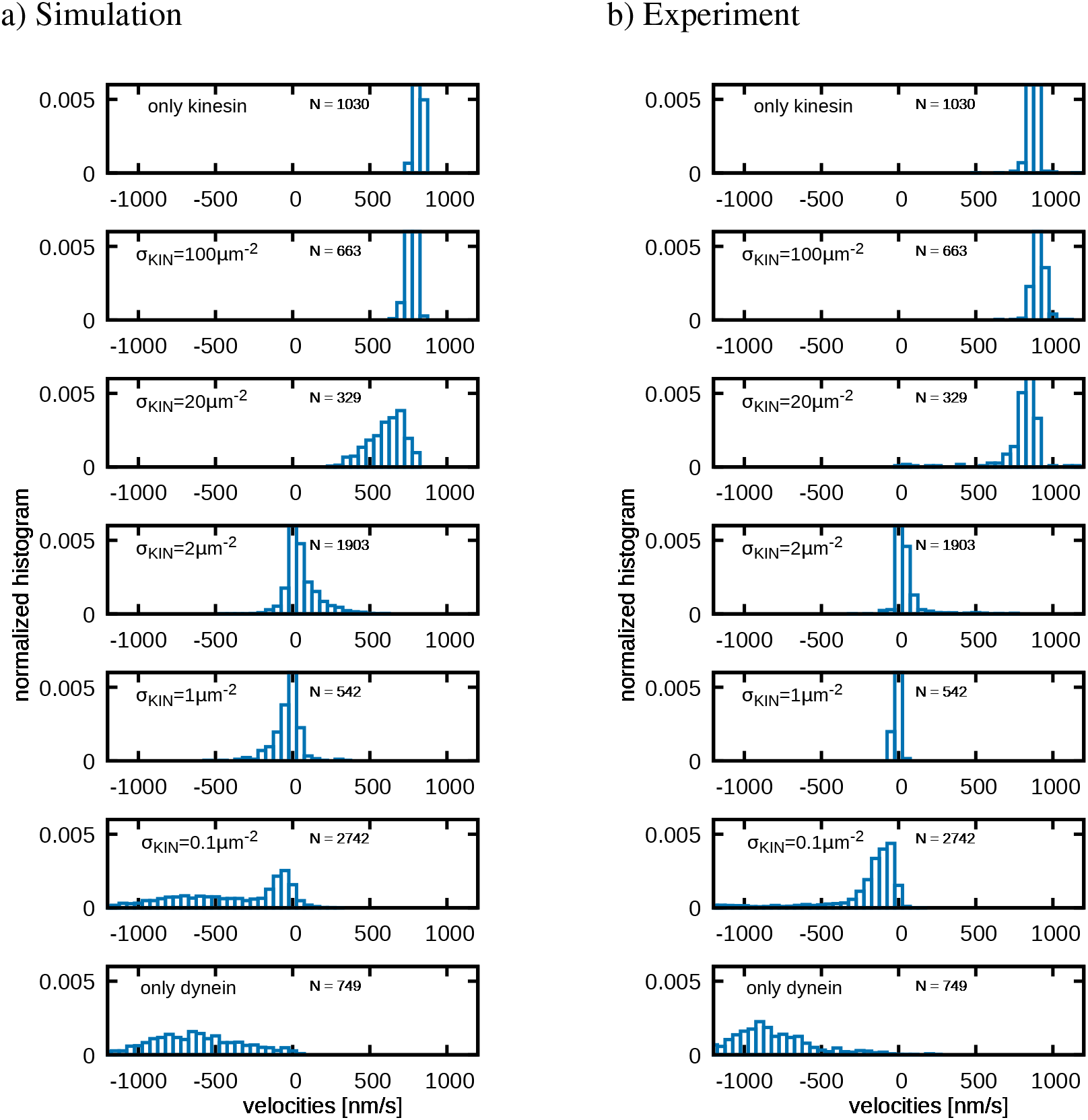
Bidirectional gliding assay simulations reproduce the experimental observations reasonably well. a) Normalized histograms of microtubule gliding velocities from simulations and b) from experiments. For simulation and experiment the same microtubule length distribution of lengths between *L*_MT_ = 10 μm and *L*_MT_ = 15 μm were applied and the dynein density was kept constant at *σ*_dyn_ = 64 μm^−2^. The kinesin density was varied as written in the plots including the “only kinesin” case with a kinesin density of *σ*_kin_ = 100 μm^−2^. With decreasing kinesin density the histograms for simulation and experiment show a transition from fast kinesin-driven transport (positive velocities) with a peak around 800 nms^−1^ towards dynein-driven transport with a wide distribution of negative velocities. For both, simulation and experiment, a balanced state is observed for kinesin densities around *σ*_kin_ = 1 – 2 μm^−2^. At higher kinesin densities (*σ*_kin_ = 20 - 100 μm^−2^) kinesin takes over in simulation and experiment and at lower kinesin density (*σ*_kin_ = 0.1 μm^−2^) dynein dominates. The number of data points N is given in the subfigures.

We observed that kinesin and dynein teams balanced each other at high dynein density (64 μm^−2^) but low kinesin density (1 μm^−2^). This rises the question why more dynein than kinesin motors are needed to balance each other. It might be that the dynein attachment rate is simply lower than the kinesin attachment rate and therefore more dynein is needed. Or it might be that dynein is not as strong as kinesin resulting in a higher number of dynein motors needed to balance kinesin. To understand the mutual interplay between kinesin and dynein motors, first the influence of different dynein densities is considered. Different dynein densities have been shown to strongly influence the transport velocity in unidirectional dynein transport (Monzon and Scharrel et al., 2019). However, the influence of the number of dynein motors on bidirectional transport has not been studied. Here, we measured for different constant dynein densities the median microtubule gliding velocity as a function of the kinesin density for simulation (Fig. 4a) and experiment (Fig. 4b). For intermediate (13 μm^−2^) and high (64 μm^−2^ and 128 μm^−2^ for simulation only) dynein densities, a transition from the dynein-driven (negative velocities) to the kinesin-driven state (positive velocities) was observed with increasing kinesin density. The lower the dynein density the more the balance was shifted to lower kinesin densities. At the lowest dynein density (3 μm^−2^) no dynein-driven state was observed but rather the balanced state extended to low kinesin densities (see Fig. 4a). But a kinesin-driven state was observable. In conclusion these finding’s show that besides the kinesin density, the dynein density also regulates the directionality of bidirectional transport.

**Fig. 4.**
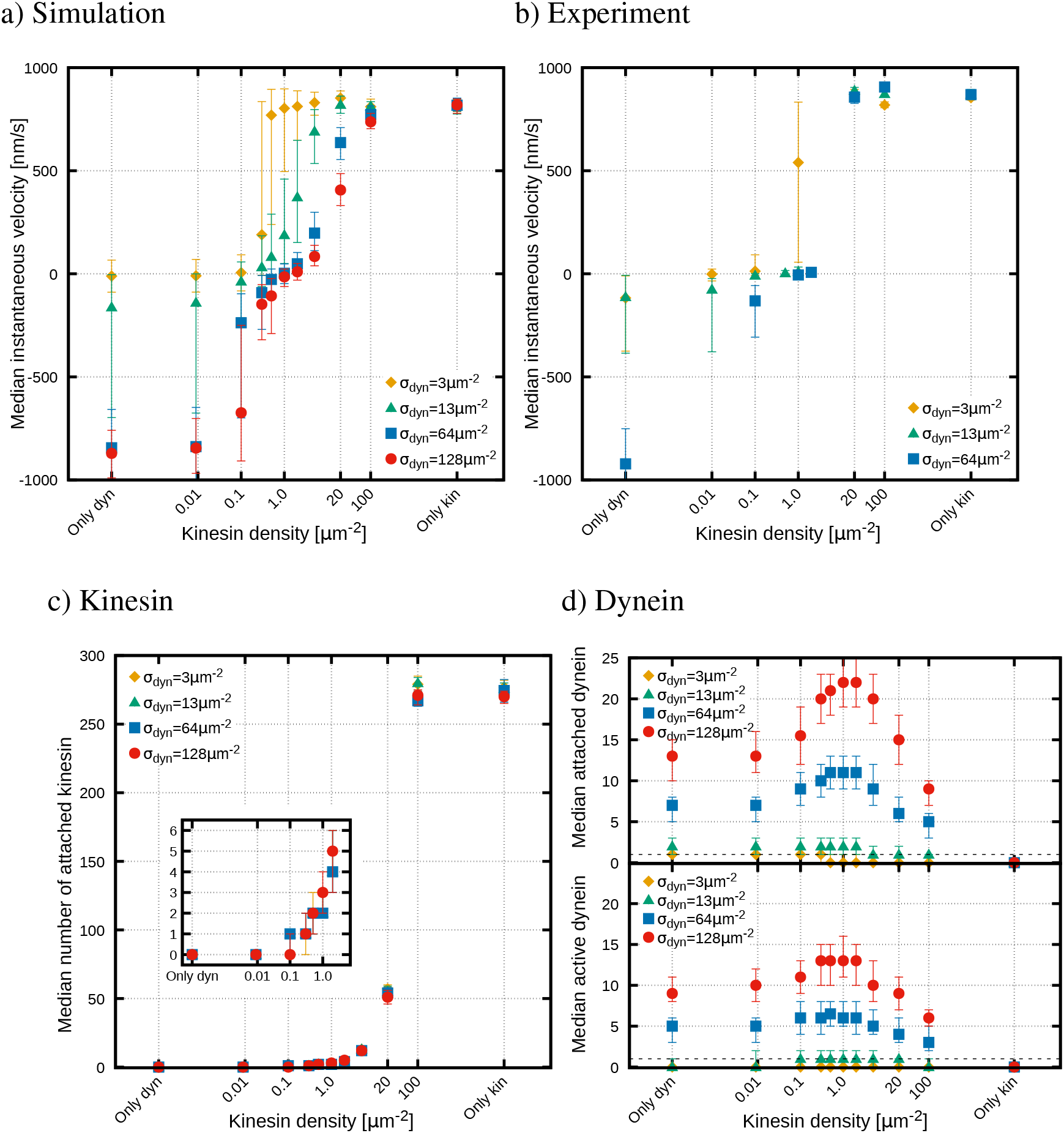
Dynein density regulates bidirectional transport and kinesin stabilizes the balanced state by activating passive dynein and increasing the number of attached dynein. a) Bidirectional gliding assay simulations at different constant dynein densities. Median instantaneous velocities with interquantil range (IQR) are shown as a function of varying kinesin densities. The microtubule length is 25 μm. For all dynein densities we see a kinesin-driven state for *σ*_kin_ ≥ 20 μm^−2^ and for all dynein densities, except *σ*_dyn_ = 3 μm^−2^, a dynein-driven state for *σ*_kin_ ≤ 0.1 μm^−2^. For the lowest dynein density median velocities in the dynein driven state are zero as in the balanced state. For intermediate and high dynein densities (*σ*_dyn_ = 13, 64 and 128 μm^−2^) the balanced states shift towards lower kinesin densities the lower the dynein density is. b) Bidirectional gliding assay experiments at different constant dynein densities. Median instantaneous velocities with IQR are shown as a function of varying kinesin densities. The shown data is from microtubules with lengths *L*_MT_ > 12 μm. For all dynein densities we see a kinesin-driven state for *σ*_kin_ ≥ 20 μm^−2^ and a dynein-driven state for very low kinesin densities (*σ*_kin_ ≤ 0.01 μm^−2^ for intermediate dynein density and *σ*_kin_ ≤ 0.1 μm^−2^ for the highest dynein density). In agreement with the simulation (Fig. 4a) the balanced state shifts towards lower kinesin densities the lower the dynein density is. c) Median number of attached kinesin as a function of varying kinesin densities at different constant dynein densities corresponding to a). The number of attached kinesin increases monotonically with increasing kinesin density and does not significantly depend on the dynein density. d) Median number of total (active and passive) attached dynein (upper panel) and median number of active attached dynein motors (lower panel) corresponding to a). Median numbers are depicted as a function of the varying kinesin density for different constant dynein densities. The horizontal dashed line represents the one. For the lowest dynein density the total attached dynein number is one in the dynein-driven state and decreases to zero as soon as a kinesin is attached (*σ*_kin_ ≥ 0.3 μm^−2^, see Fig. 4c). However, the attached dynein never activates (see lower panel for median active dynein number). For the intermediate dynein density two dynein motors are attached in the dynein-driven and the balanced state. As soon as a kinesin is attached (*σ*_kin_ ≥ 0.1 μm^−2^, see 4c) one of the two attached dynein motors is active. For high dynein densities (*σ*_dyn_ = 64 μm^−2^ and *σ*_dyn_ = 128 μm^−2^) the total attached dynein number and the active attached dynein number reach their maxima at the kinesin density of the balanced state (see Fig. 4a).

As stated above, at the lowest dynein density (3 μm^−2^) the balanced and the dynein-driven state could not be distinguished from each other since the velocity was close to zero also for the dynein-driven state. This means for the dynein-driven state at the lowest dynein density two scenarios are possible: (i) either no dynein is attached to the microtubules or (ii) the attached dynein does not move pointedly towards the microtubule minus end. To understand which scenario is at play, the median number of attached dynein motors was determined as function of the kinesin density using the simulation (Fig. 4d, upper panel). We see that one dynein motor is attached for the lowest dynein density and kinesin densities smaller than 0.5 μm^−2^. This rules out the first possibility that no dynein motor is attached at all. To test whether the second scenario holds, we need to look at the internal dynein state (passive or active). Unlike kinesin, which is always active in the model, dynein is known to have an active and a passive state (Zhang et al., 2017; Torisawa et al., 2014; Monzon and Scharrel et al., 2019). In our model processive motion towards the microtubule minus-end is performed by active dynein. Whereas diffusion in a harmonic potential is implemented for attached passive dynein (See Material and Methods for details of the model)^3^. Therefore, passive attached dynein motors does not exert directed forces on the microtubule. To exert directed forces, passive attached dynein motors need to activate. Here, a mechanical dynein activation is modeled, *i.e*. mechanical stretching activates passive attached dynein. A passive attached dynein motor is stretched when the microtubule is transported by other motors. This means, one attached passive dynein motor alone will never activate. Consequently, in the “only dynein” case the one attached dynein motor will never activate and, therefore, never walks processively towards the microtubule minus-end. Thus, no net-movement occurs. However, as soon as a kinesin motor is attached, the kinesin motor could transport the microtubule and activate the passive attached dynein motor. To know whether an attached kinesin activates the one attached passive dynein, the number of attached active dynein motors (Fig. 4d, lower panel) as well as the number of attached kinesin (Fig. 4c) was determined using the simulation. We see the median number of active dynein is always zero for the lowest dynein density. Moreover, we see that as soon as one kinesin was attached, no dynein was attached anymore for the lowest dynein density. This means kinesin directly pulled off the one attached dynein motor. As a consequence, a balanced state with balanced forces between kinesin and active dynein does not exist. Thus, one dynein cannot resist against kinesin.

There are two possible reasons why one dynein cannot resist against kinesin: First dynein is passive and therefore cannot resist against kinesin or second dynein is activated for a short moment but then directly pulled off by kinesin since it is not strong enough. To understand what kind and how many dyneins are needed to resist against kinesin, we look at a slightly higher dynein density. For the intermediate dynein density (13 μm^−2^) we observed a clearly separated balanced state (Fig. 4a and 4b). This means for intermediate dynein density a force balance state where dynein resist against kinesin exists. Looking again at the number and internal states of attached motors (Fig. 4c and 4d), we see two dynein motors are attached in the balanced and the dynein-driven state. While in the dynein-driven state the median number of active attached dynein is zero^4^, in the balanced state one of the two dynein motors is active. Thus we see as soon as one kinesin is attached, the kinesin activates one of the two passive attached dynein motors. This means for the lowest dynein density kinesin activated dynein and then directly pulled it off. This implies that one active dynein motor is not strong enough to resist against one kinesin motor. A previous study defined a motor as “strong” when having a large “stall force to detachment force ratio” (*F*_s_/*F*_d_) and as “weak” when having a small “stall force to detachment force ratio” (*F*_s_/*F*_d_) (Müller et al., 2008). To determine if kinesin and dynein are strong or weak motors we compare kinesin and dynein single molecule parameters (see Fig. 2d and 2e for the modeled parameters and the parameter list for references). Kinesins detachment force is as high as its stall force (6pN) resulting in a “stall force to detachment force ratio” of exactly one. The stall force of dynein is 1.25 pN. Unlike kinesin, which has a symmetrical detachment behavior under forward and backward load, dynein detaches faster under forward loads (assisting forces) than under backward loads (resisting forces) (Fig. 2d and 2e). The dynein detachment force is 4 pN under backward load and 0.125 pN under forward load. Thus in the tug-of-war situation (mainly backward load) the “stall force to detachment force ratio” of dynein is 0.3125. Therefore, dynein is the weaker motor in the competition with kinesin. That is why, one active dynein is directly pulled off by kinesin. However, we observed that for the intermediate dynein density one active dynein could resist against kinesin. The reason is that at the intermediate dynein density additionally to one active dynein a passive dynein was attached. If, in this case, the active dynein is pulled off by kinesin, the passive dynein helps out. Then the passive dynein is activated by kinesin and tries to resist against kinesin until it is also pulled off. Importantly, in the mean time a new passive dynein can attach and serve again as a substitute for the new active dynein. We conclude the role of the passive dynein is helping out when active dynein is pulled off by kinesin. Consequently, an active dynein having a passive dynein as a substitute is able to temporally hold against one kinesin. Whereas one active dynein alone cannot resist against kinesin due to being the weaker motor in the competition with kinesin.

Our findings showed that two dynein can temporally hold against one kinesin. However, the balanced state of two dynein motors competing against one kinesin is not stable, because the active dynein is continuously pulled off. Once a new passive dynein does not attach fast enough to help the new active dynein, kinesin will take over completely. Obviously, a higher number of dynein motors are expected to stabilize the balanced state. At high dynein densities of 64 μm^−2^ (same for 128 μm^−2^ in simulation) a stable balanced state with only small fluctuations was observed (Fig. 4a and 4b). Here, two kinesin motors competed against 6 active dynein motors in the balanced state (Fig. 4c and 4d). Thus indeed, more active dynein motors were attached. Furthermore, the number of active dynein motors was maximal in the balanced state. This means kinesin activated more dynein and thereby increased the number of active dynein. Kinesin, therefore, stabilized the balance state. Moreover, besides the number of active dynein, the total number of attached dynein (active and passive) reached its maximum in the balanced state. This means besides activating dynein the activity of kinesin reduced the overall dynein detachment. Having kinesin pulling back dynein and thereby reducing the dynein detachment could be interpreted as an effect coming from a dynein catch-bond. However, rather than a catch-bond, a linearly increasing detachment rate is implemented for single dynein. Therefore, the reason has to be a collective effect. Imagine having some dynein and a few kinesin motors attached. Attached kinesin motors transport the microtubule in a kinesin direction. Thereby kinesin aligns attached dynein motors under backward load (see Fig. 2c for an illustration). Under backward load the dynein detachment increases more slowly with the force than under forward load (see Fig. 2e, lower panel). Consequently, by aligning the dynein motors under backward load, less dynein motors detach compared to the “only dynein” case, where the deflections of attached dynein motors are in random directions (see Fig. 2b and 2c for an illustration)^5^. In the “only dynein” case some motors might be under backward load, others under forward load resulting in a higher overall detachment rate. In conclusion, kinesin stabilizes the balanced state first by activating passive dynein and second by reducing the dynein detachment when aligning dynein motors under backward load.

In summary, single dynein is a weaker motor. A single dynein would not be able to transport a cargo. However, in a team the functionality of dynein is increased. In a team, passive dynein helps out when active dynein is pulled off. Additionally, dynein is strengthened by the activity of the stronger kinesin. The activity of kinesin on one hand activates passive attached dynein and on the other hand increases the number of attached dynein. Because kinesin increases the number of attached dynein and because weaker active dynein needs passive dynein as a substitute, the number of dynein motors is higher than the number of kinesin motors in the balanced state. In total this means while one kinesin motor is strong by itself (see also 4c to see that the number of kinesin motors does not significantly change with the dynein density), the dynein strength depends on other dynein and kinesin motors. However, we showed the directionality of bidirectional transport can be regulated by varying the number of kinesin or the number of dynein motors.

### Stable balanced state upon different ATP concentrations

Single molecule velocities of dynein and kinesin strongly depend on ATP concentration (Schnitzer et al., 2000; Ross et al, 2006; Torisawa et al., 2014). However, it is unclear whether the directionality of bidirectional transport can be regulated by changing ATP concentration. To understand the influence of ATP concentration on bidirectional transport in gliding assays, it is important to know how individual teams of kinesin and dynein motors are influenced by ATP concentration. Therefore, first median velocities of unidirectional gliding assays as a function of ATP concentration were measured (Fig. 5a and 5b). In both, kinesin and dynein unidirectional gliding assays an intermediate motor density (18 μm^−2^) was used. We observed median velocities of the kinesin and the dynein assay increased with increasing ATP concentration for simulation and experiment. While median velocities of the kinesin assay could perfectly be fitted with a Michaelis-Menten equation, velocities of the dynein assay did not show such a behavior. In the dynein assay the velocity increased more in a linear manner and did not saturate at the highest applied ATP concentrations. This means at the highest applied ATP concentrations, kinesin velocity does not change with changing ATP concentration anymore, while the dynein velocity does. This means kinesin and dynein, indeed, react differently to ATP concentration. Changing the ATP concentration, therefore, might regulate the directionality of bidirectional transport.

**Fig. 5.**
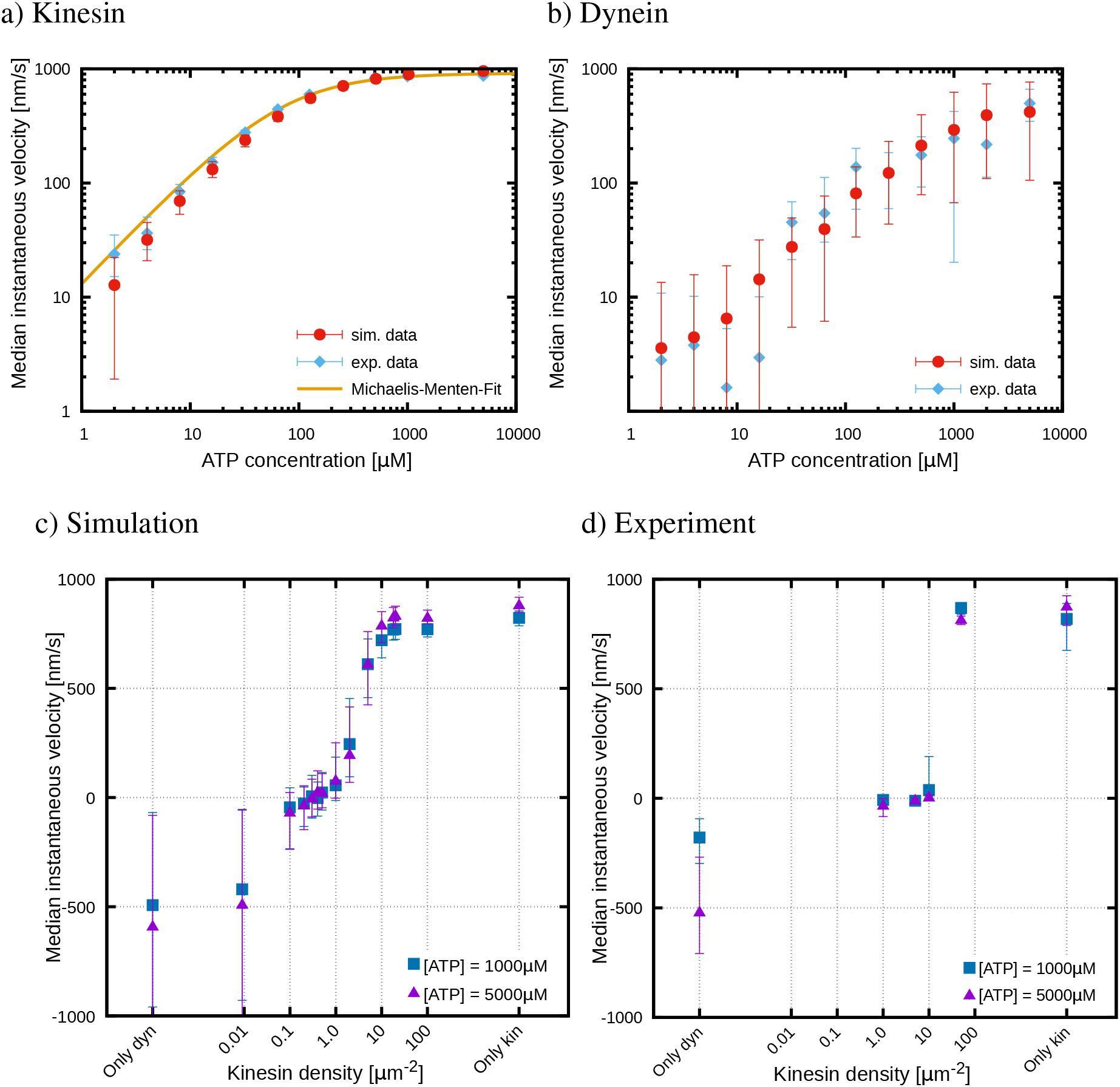
Although unidirectional kinesin and dynein assays react differently to ATP concentration, changes of ATP concentration cannot regulate the direction of bidirectional transport. a) Median instantaneous gliding velocities (with IQR) as a function of ATP concentration for the unidirectional kinesin assay. For all ATP concentration a constant kinesin density of *σ*_kin_ = 18 μm^−2^ was applied in simulation (red curve) and experiment (light blue curve). Experiment and simulation results show a perfectly matching Michaelis-Menten dependence on the ATP concentration. To the experimental data a Michaelis Menten equation (*v* = *v*_max_ × [ATP]/(*K_m_* + [ATP]) with *v*_max_ = 914 nm s^−1^ and *K_m_* = 69 μM was fitted (solid fit). b) Median instantaneous gliding velocities with IQR as a function of ATP concentration for the unidirectional dynein assay. For all ATP concentration a constant dynein density of *σ*_kin_ = 18 μm^−2^ was applied in simulation (red curve) and experiment (light blue curve). In simulation and experiment the same microtubule length distribution of microtubules with lengths bigger than 15 μm was used. Experimental and simulation results perfectly match but could not befitted with a meaningful Michaelis-Menten equation. Median instantaneous gliding velocities increase more in a linear manner with the ATP concentration and might not be saturated at the highest measured ATP concentration. c) Bidirectional gliding assay simulations at different ATP concentrations. Median instantaneous velocities with IQR are presented as a function of the varying kinesin density and a constant dynein density of *σ*_dyn_ = 18 μm^−2^. The microtubule length was *L*_MT_ = 25 μm. We see that, like in the experiment (Fig. 5d) the dynein-driven region is stronger influenced by ATP concentration than the kinesin-driven region. However, the balanced state remains at the same kinesin density (*σ*_kin_ = 0.1 - 1.0 μm^−2^) for all ATP concentrations. d) Bidirectional gliding assay experiments at different ATP concentrations. Median instantaneous velocities with IQR are presented as a function of the varying kinesin density and a constant dynein density of approximately *σ*_dyn_ = 18 μm^−2^. The shown data is from microtubules with lengths *L*_MT_ > 15 μm. We see, like in the simulation (Fig. 5c), the velocities in the dynein-driven state are stronger reduced by lower ATP concentrations than in the kinesin-driven state. The balanced state, however, remains at the same kinesin density (*σ*_kin_ = 5 μm^−2^) for all tested ATP concentrations.

To investigate the potential regulation of bidirectional transport by ATP concentration, we applied two high ATP concentrations, where the velocity of kinesin is constant, while the velocity of dynein doubles. Therefore, we used a constant intermediate dynein density (same as for the unidirectional assay) and varied the kinesin density at two different ATP concentrations for simulation (Fig. 5c) and experiment (Fig. 5d). For both ATP concentrations, we observed the three formerly described motility states. If ATP concentration would regulate bidirectional transport, the balanced state would be expected to shift towards lower or higher kinesin densities depending on ATP concentration. However, we found the balanced state stayed at the same kinesin density for all tested ATP concentrations. This means the balanced state is invariant to ATP concentrations.

To understand why the balanced state does not depend on ATP concentration, we used again the simulation for an insight to events at the molecular level. In the balanced state, we observed mainly diffusive stepping of passive dynein (Fig. S1). Diffusive dynein stepping also depends on ATP concentration, but passive dynein does not exert directed forces on the microtubule. Therefore, changing the diffusive stepping of passive dynein does not shift the force balance to neither site. Only directional stepping of active motors could change the force balance. But in the balanced state active dynein and kinesin almost do not move at all (Fig. S1). Consequently, ATP concentration cannot shift the force balance between kinesin and dynein. Thus, the balanced state is stable upon changing ATP concentration as long as the applied forces do not depend on the ATP concentration (see Fig. S2 for more ATP concentrations). This means only the kinesin and dynein densities determine the motility state, but not the ATP concentration. In conclusion, ATP concentration cannot regulate the directionality of bidirectional transport.

### Stable balanced state in the presence of roadblocks

Previous work suggested roadblocks might regulate bidirectional transport by asymmetrically inhibiting the motor activity (Siahaan et al., 2019; Tan et al., 2019; Monroy et al., 2018). Single kinesin and dynein motors indeed show different behavior upon encountering a roadblock (Telley et al., 2009; Dixit et al., 2008; Schneider et al., 2015). On one hand single kinesin is known to either pause when encountering a roadblock or completely detach from the microtubule (Ferro et al., 2019; Dixit et al., 2008; Schneider et al., 2015). On the other hand dynein is known for frequently taking side steps (Wang et al., 1995) and therefore able to circumvent roadblocks without detaching. Single dynein has been shown to overcome roadblocks more successfully than single kinesin (Ferro et al., 2019). However, in cargo transport by multiple motors, multiple kinesins and multiple dyneins appear to be affected by roadblocks in a similar manner (Ferro et al., 2019). It is therefore of interest, to study if roadblocks alter the transport behavior in bidirectional gliding assays.

To test if dynein is as affected by roadblocks as kinesin also in unidirectional gliding assays, we performed unidirectional kinesin and dynein gliding assays at different roadblock concentrations. Therefore, we coated the microtubule with rigor binding kinesin-mutants (hereafter referred to as roadblocks) (Schneider et al., 2015) at different concentrations. For different roadblock concentrations, we measured median gliding velocities at a constant dynein or kinesin density of 50μm^−2^, respectively (see Fig. 6a simulation and 6b experiment). In both gliding assays, the median velocity decreased with increasing roadblock concentration for simulation^6^ and experiment. Comparing the results of the kinesin and dynein gliding assays, we find that dynein is more affected than kinesin for experiment and simulation. This is in disagreement with our expectations from previous studies stating that kinesin is as affected as dynein (Ferro et al., 2019). Since dynein likely uses side steps as its bypassing mechanism (Reck-Peterson et al., 2006), we hypothesise that dynein might not be able to take side steps in the gliding assay set-up. To test our hypothesis we used the simulation to implement several protofilaments and allowed dynein to take side steps. Using the several-protofilament simulation, we observed that dynein was as affected by roadblocks as kinesin (Fig. S3a). Our findings from the several-protofilament simulations agree with findings of Ferro et al. (2019) for cargo transport. However, not allowing dynein to take side steps in the multiple protofilament simulation, dynein is again more affected than kinesin. Consequently, if dynein does not take side-steps, both motors have to detach and reattach to overcome roadblocks. Comparing dynein and kinesin attachment rates (Fig. 2b and 2c), we see the kinesin attachment rate is much higher than the dynein attachment rate. Therefore, kinesin can reattach a lot faster than dynein. We conclude that in microtubule gliding assays dynein is not able to take side steps. Without its side stepping ability, dynein is more affected by roadblocks than kinesin.

**Fig. 6.**
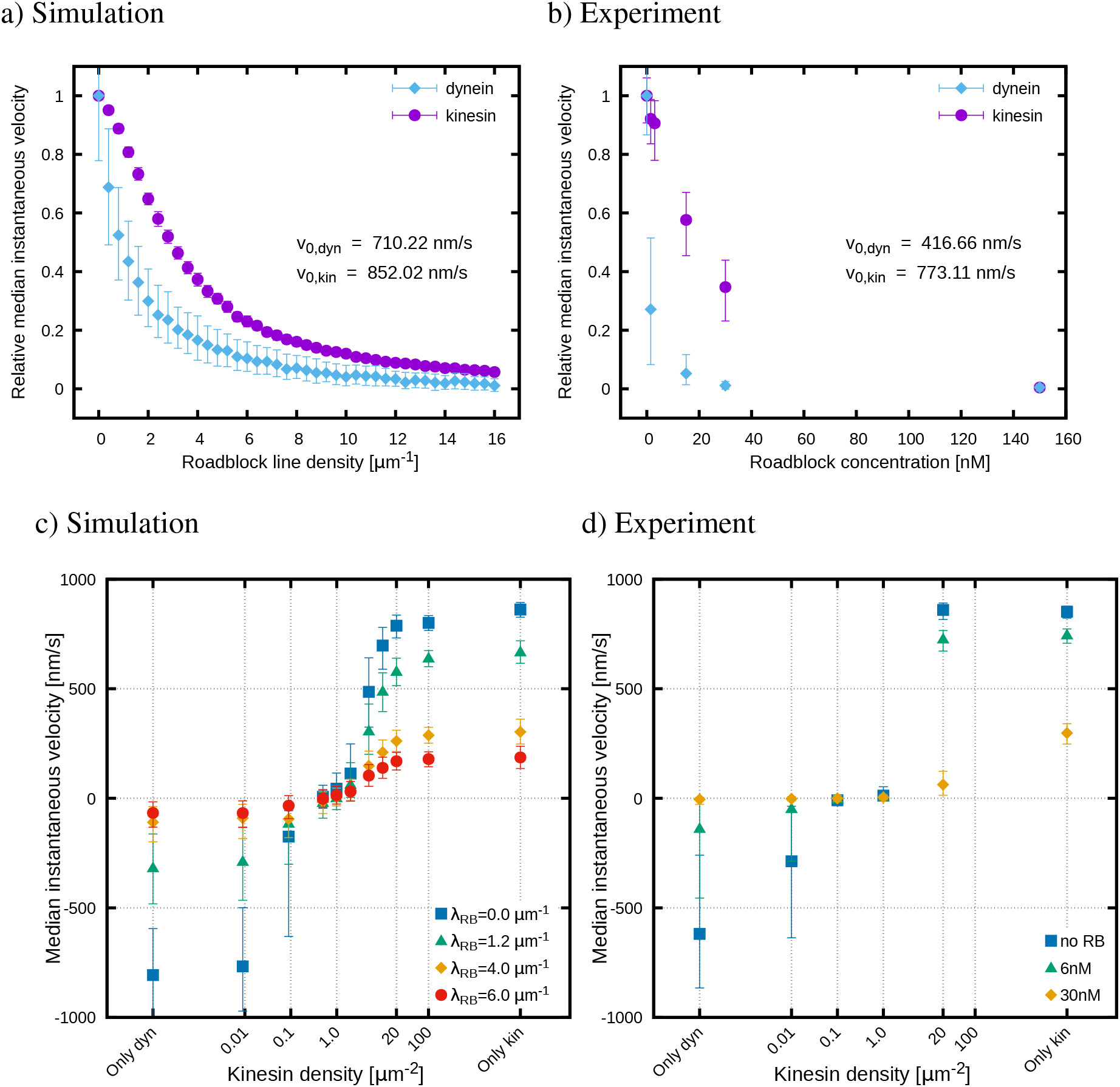
Although unidirectional kinesin and dynein assay are differently affected by roadblocks, roadblocks cannot change the transport direction of bidirectional transport. In this set-up the microtubule is coated with rigor binding kinesin mutants referred to as roadblocks. While in the experiment a roadblock concentration is given, in the simulation a roadblock line density λ_RB_ was applied. a) Unidirectional kinesin (purple curve) and dynein (light blue curve) assay simulations in the presence of roadblocks. Median instantaneous velocities with IQR relative to the velocity in the absence of roadblocks are depicted as a function of the roadblock line density. In both, the kinesin and the dynein assay, a microtubule length of *L*_MT_ = 25 μm and motor densities of *σ*_kin_ = *σ*_dyn_ = 50 μm^−2^ were used. In both assays the velocity decreases with increasing roadblock line density. In agreement with the experiment (Fig. 6b) the gliding velocity of the dynein assay decreases faster with the roadblock line density than the velocity in the kinesin assay. b) Experiments of unidirectional kinesin and dynein assays. Median instantaneous velocities with IQR relative to the velocity in the absence of roadblocks are depicted as a function of the applied roadblock concentration. In both, the kinesin and the dynein assay, microtubule lengths of the intervall *L*_MT_ = 25 - 30 μm and motor densities of *σ*_kin_ = *σ*_dyn_ = 50 μm^−2^ were used. As in the simulation for both assays the velocity decreases with increasing roadblock concentration, whereby velocities of the dynein assay decrease faster than of the kinesin assay. c) Bidirectional gliding assay simulations at different roadblock line densities. Median instantaneous velocities with IQR are depicted as a function of varying kinesin densities. The dynein density was kept constant at *σ*_dyn_ = 50 μm^−2^ and the microtubule length was *L*_MT_ = 25 μm. We see the dynein-driven state is stronger affected by roadblocks than the kinesin-driven state. However, the balanced state remains at the same kinesin density of *σ*_kin_ = 1.0 μm^−2^ for all roadblock line densities. d) Bidirectional gliding assay experiments at different roadblock concentrations. Median instantaneous velocities with IQR are shown as a function of varying kinesin densities at a constant dynein density of *σ*_dyn_ = 50 μm^−2^. In agreement with the simulation (Fig. 6c), the dynein-driven state is stronger affected by roadblocks than the kinesin-driven state. But the balanced state remains at the same kinesin density of *σ*_kin_ = 0.1 - 1.0 μm^−2^ for all roadblock concentrations.

We observed that multiple dynein and kinesin motors react differently to roadblocks in microtubule gliding assays. This rises the question if the different reaction to roadblocks influences bidirectional transport in gliding assays. Therefore, we applied the same constant dynein density as for the unidirectional assay with roadblocks and varied the kinesin densities at different roadblock concentrations. In the simulation for all roadblock densities^6^, clear motility states were observable (Fig. 6c). In the experiment the dynein-driven state merged with the balanced state at very high roadblock concentrations (Fig. 6d). However, for simulation and experiment the balanced state stayed at the same kinesin density for all roadblock concentrations. Additionally, we also implemented several protofilaments with dynein taking side steps. For the bidirectional gliding assay simulation with several protofilaments, the balanced state also stayed at the same kinesin density for all roadblock concentrations (Fig. S3b). As a consequence, independently of the side stepping ability of dynein, the balanced state remains at the same kinesin density for all roadblock concentrations.

To obtain an explanation for the balanced state being stable in the presence of roadblocks, we used again the simulation for insights at the molecular level. As stated before, kinesin and active dynein almost did not move at all in the balanced state. Here, we measured in detail the moved distance of kinesin and dynein in the balanced state (Fig. S4). The median moved distance is 65 nm for kinesin and 16 nm for dynein. Compared to the mean distance between roadblocks (approximately 166.67 nm for a roadblock density of 6 μm^−1^), the moved distance of kinesin and dynein in the balanced state is small. Thus, the motors are strongly localized in the balanced state. Because of the strong motor localization, the balanced state is stable in the presence of roadblocks. Consequently, the kinesin and dynein densities determine the motility state independently of the presence of roadblocks. In conclusion, roadblocks do not regulate the directionality of bidirectional transport.

Taken together, we observed that ATP concentration and roadblocks can alter the velocities in the kinesin- or dynein-driven states, but the balanced state remains stable upon different ATP and roadblock concentrations. Exclusively, the number of motors regulates the tug-of-war between opposite-directed teams of kinesin and dynein motors.

## DISCUSSION

In this work we showed that neither ATP nor roadblock concentrations regulate the directionality of bidirectional transport. While we have seen that ATP and roadblock concentrations influence the dynein- and kinesin-driven states, the balanced state remains stable. In the balanced state, active dynein and kinesin almost do not step at all. This suggests that parameters influencing the stepping of the motors cannot be used to change the balanced state and therefore cannot regulate the directionality of bidirectional transport. Consequently, the inability of dynein to take side steps in our bidirectional gliding assays does likely not change our result, i.e. even if dynein was able to take side steps during cargo transport, roadblocks would also not influence the balanced state. Unlike ATP and roadblock concentrations, the number of motors, indeed, regulates the directionality of bidirectional transport. By changing the number of involved motors, the balanced state can be shifted towards a kinesin- or dynein-driven state. The number of motors influences the force balance between kinesin and dynein motors. Having more kinesin motors, the kinesin team is stronger and vice versa. Therefore, factors that influence the force balance between the kinesin and dynein team should play a key role in the regulation of bidirectional transport.

Our results show that kinesin activity influences the force balance. Using a mathematical model, we showed that the kinesin activity strengthens dynein. The kinesin activity on one hand activates passive attached dynein motors and on the other hand increases the number of total attached dynein. As a first consequence, the number of attached dynein motors in the balanced state is higher than the number of kinesin motors. This is in agreement with previous studies reporting that more dynein motors than kinesin motors are involved in the tug-of-war (Soppina et al., 2009; Hendricks et al., 2010). A second consequence is that kinesin stabilizes the balanced state by strengthening the weaker and partly passive dynein motors. This explains why we find a stable force balance between kinesin and dynein. Thus, in addition to antagonistic effects between kinesin and dynein teams, there are also cooperative effects. The cooperative effect of kinesin stabilizing the balance state could be used to hold a cargo at a specific location inside the cell as it was hypothesized by Encalada et al. (2011) for PrPC vesicles.

We find that from the three possible regulation factors, number of motors, ATP concentration and roadblocks, only the number of motors could regulate bidirectional transport. However, varying the number of motors might not be the only way the stable force balance can be regulated. Adding adaptor proteins that activate passive attached dynein could likely also change the force balance. We showed that one single mechanically activated dynein without adaptor proteins is pulled off directly by kinesin. This is in agreement with the work of Belyy et al. (2016), where they showed that one single dynein cannot hold against a single kinesin. However, Belyy et al. (2016) showed that a single dynein with adaptor proteins (dynein-dynactin-BicD2 complex) can hold against a single kinesin. Moreover, measuring dynein stall force, they found out that dynein plus adaptor proteins has a stall force of 4. 4 pN. This is around three to four times more than the stall force of a single dynein without adaptor proteins (1.0 – 1.25 pN) (Mallik et al., 2004). Thus, dynein activated by adaptor proteins is stronger than mechanically activated dynein motors, and adaptor proteins should be able to shift the force balance. Based on our work showing that factors influencing the force balance are key regulators of bidirectional transport, we would predict that adaptor proteins should be able to regulate bidirectional transport by increasing the dynein stall force. This might suggest that even kinesin-driven transport could be reversed by adding adaptor proteins.

Beside adaptor proteins changing the dynein stall force and therefore affecting the force balance, it has been shown that the ATP concentration can also change the dynein stall force. Mallik et al. (2004) showed that for ATP concentrations lower than 1000 μM, the dynein stall force increases linearly with ATP concentration. For ATP concentrations above 1000 μM, the stall force does not depend on ATP concentration. In our work, we showed that for ATP concentrations equal to or greater than 1000 μM, the ATP concentration does not influence the force balance. However, the force balance could be influenced by lower ATP concentrations. Simulating the bidirectional gliding assay at lower ATP concentration, we did in fact see a tendency of the balanced state to be moved towards lower kinesin densities (Fig. S2b). But the effect was as small as shifts arising from fluctuations in the number of motors involved in the transport. However, in Klein et al. (2014) the transport direction could be significantly changed by ATP concentration in cargo transport simulations. Unlike in microtubule gliding assays, in cargo transport simulations, the number of involved kinesin and dynein motors is constant. That is why fluctuations are smaller and the effect of the ATP concentration altering the force balance is visible. Thus, we predict that in microtubule gliding assays shifts of the force balance stemming from altered ATP concentrations are small compared to shifts stemming from fluctuating numbers of motors.

An additional regulation factor influencing the force balance could be roadblocks that act exclusively on just one kind of motor. This study shows that the presence of roadblocks hindering both motors does not change the force balance. However, having roadblocks which asymmetrically influence the motors could have an influence. Tau islands, for instance, have an asymmetric effect on motor proteins. While kinesin detaches when encountering a tau island, dynein continues walking (Tan et al., 2019; Siahaan et al., 2019). This means the tau island changes the motor number and therefore the force balance. Having a kinesin-driven transport which encounters a tau island, for instance, would cause a shift to a dynein-driven state because kinesin motors would detach and dynein could take over. Therefore, roadblocks which asymmetrically detach one kind of motor^7^ can be understood to alter the number of attached motors and therefore shift the force balance. As a consequence, such roadblocks could be used to regulate bidirectional transport.

Our finding that dynein gliding assays do not show a Michaelis-Menten-like dependence on ATP concentration is in contrast to previous work of Torisawa et al. (2014). To understand why we do not see a Michaelis-Menten-like ATP dependence, we simulated the unidirectional gliding assay at a higher dynein density and varying ATP concentrations. At a higher dynein density (Fig. S2a) the simulation indeed shows such a dependence on ATP concentration. We know from our previous work (Monzon and Scharrel et al., 2019) that at low dynein densities, passive dynein motors slow down the microtubule gliding (Note that diffusive stepping of passive dynein also increases with ATP concentration, see Material and Methods for the model details). This means the lower the ATP concentration the slower the passive dynein motors step and the more the passive motors slow down the microtubule gliding. This explains why the increase in gliding velocity with the ATP concentration at lower dynein density is slower than predicted by a Michaelis-Menten equation. Consequently, we do not see a Michealis-Menten-like dependence on ATP concentration at low and intermediate dynein densities. At high dynein density, the slowing down by passive dynein motors is negligible and we see a Michaelis-Menten dependence on ATP concentration. This implies the ATP dependence of dynein-driven transport depends on the number of dynein motors.

In our study, we have provided key insights into the factors influencing bidirectional transport in the absence of adaptor proteins. Here, the factors that influence the force balance seem to be most important, whereas others such as stepping or bypassing mechanisms have little effect. Future work should now focus on the influence of adaptor proteins in light of this or roadblocks differentially influencing the number of attached motors.

## MATERIAL AND METHODS

### Materials and reagents

All reagents unless otherwise stated were purchased from Sigma. cDNA encoding full length D.melanogaster Kinesin Heavy Chain (KHC) was sub-cloned into an inhouse generated insect cell expression vector in frame with a C-terminal 6xHis tag. Codon optimized gene sequence of R.norvegicus kif5c (GenArt, gene synthesis, Invitrogen) truncated to first 430 amino acids with a T39N rigor mutation was cloned upstream of 6xHis affinity (with and without eGFP). Plasmids encoding genes for H.sapiens dynein subunits were a kind gift from Max Schlager and Andrew Carter.

### Microtubules

Tubulin was purified from pig brains by two cycles of polymerization and depolymerization in high concentration PIPES buffer (Castoldi et al., 2003), labelled with Alex 488 dye, diluted to 4 mg/mL, aliquoted and stored at −80° C.

Polarity marked microtubules were prepared by preferentially adding a short brightly labelled seed to plus ends of GMPCPP and taxol polymerized microtubules (referred to as ‘double stabilized’ microtubules) in the presence of N-Ethylmaleimide (NEM)- labelled tubulin. This was carried out as follows: 4.8 μM tubulin was freshly labelled with 1 mM NEM in the presence of 0.5 mM GTP for 20 min on ice. Excess NEM reagent was quenched with 8 mM β-mercaptoethanol. An elongation mix consisting of 1 mM GTP, 4 mM MgCl2, 1 mM Alexa-488-labelled tubulin and 0.64 μM NEM-tubulin in BRB80 was incubated on ice for 5 min, shifted to 37°C for 40 s followed by the addition of 250 μM double stabilized microtubules and incubated at 37° C for 1 hr. Resultant microtubules were stabilized with 20 μM taxol. Polarity marked microtubules were always prepared fresh every day at the beginning of an experiment.

### Motor proteins

#### Expression and purification of D.melanogaster Kinesin Heavy Chain (KHC)

KHC was purified from SF9 cells as described in (Korten et al., 2016). Briefly frozen cell pellets infected with FlexiBAC baculoviral particles for 72 hr were lysed via ultra-centrifugation at 4^°^ C. Lysates were pre-clarified over a cation exchange column and the resultant elute was passed over an His-tagged IMAC column. Column-bound KHC dimers were eluted with 300 mM imidazole, desalted and snap frozen.

#### Expression and purification of H.sapiens Dynein

Recombinant cytoplasmic dynein expression and purification was performed based on a published protocol (Schlager et al., 2014). The MultiBac plasmid was a generous gift by Max Schlager and Andrew Carter.

The MultiBac plasmid was integrated into the baculoviral genome of DH10EMBacY cells by using Tn7 transposition. 50 ng of MultiBac plasmid was incubated with 100 μL of chemical competent DH10EMBacY cells for 45 min on ice. The mix was heat shocked for 45 s at 42° C and incubated with 600 μL LB medium for 6 h at 37°C. 150 μL cell suspension was streaked out on agar plates containing 50 μg ml^−1^ Kanamycin, 10 μgml^−1^ Gentamycin, 10 μgml^−1^ Tetracycline, 20 μgml^−1^ Xgal and 1 μg ml^−1^ IPTG and incubated at 37^°^C for 2-3 days. Plasmids from white colonies were isolated and checked for presence of all subunits via PCR. SF9 insect cells were infected with bacmids at 1:1000 virus to cell ratio and grown for 4 days. Cells were harvested at 300× g for 15 min at 4°C and pellets were snap frozen in liquid nitrogen at stored at –80°C.

For purification of recombinant dynein, a frozen pellet corresponding to 250 ml insect cell culture was thawed on ice and resuspended in lysis buffer (50 mM HEPES pH 7.4,100 mM NaCl, 1 mM DTT, 0.1 mM ATP, 10 % (v/v) glycerol) supplemented with 1x protease inhibitor cocktail (complete EDTA free, Roche) to a final volume of 25 mL. Cells were lysed in a Dounce homogenizer with 20 strokes. The lysate was clarified at 504,000 × g for 45 min at 4^°^ C and added to 3 mL equilibrated (in lysis buffer) IgG Sepharose beads in a column and incubated on a rotary mixer for 6 hr. Protein-bound beads were washed with 50 mL lysis buffer and 50 mL TEV buffer (50 mM Tris–HCl, pH 7.4, 148 mM potassium acetate, 2 mM magnesium acetate, 1 mM EGTA, 10 % (v/v) glycerol, 0.1 mM ATP, 1 mM DTT, 0.1 % Tween 20). The ZZ affinity tag was cleaved off with 25 μgmL^−1^ TEV protease in TEV buffer at 4^°^ C on a rotatory mixer overnight. After TEV cleavage, beads were removed and recombinant dynein was concentrated in a 100 KDa cut-off filter (Amicon Ultracel, Merck-Millipore) to a final volume of 500 μL. The TEV protease was removed by size-exclusion chromatography using a TSKgel G4000SWXL column equilibrated in GF150 buffer (25 mM HEPES pH 7.4, 150 mM KCl, 1 mM MgCl2, 5 mM DTT, 0.1 mM ATP, 0.1 % Tween 20). Peak fractions were collected, pooled and concentrated to a maximum concentration of 1 mg/ml with 100 KDa cut-off filter. All purification steps were performed at 4°C. Recombinant dynein was frozen in liquid nitrogen in the presence of approximately 10 % (v/v) glycerol. Protein concentration was determined with Bradford reagent.

#### Expression and purification of rigor rkin430

A preculture of rkin430-T39N transformed E.coli BL21 pRARE cells was grown overnight in LB medium at 37° C. 750 mL fresh LB medium, supplemented with 50 μg/mL Kanamycin, was inoculated with 5 mL preculture and incubated in a shaker at 180 rpm, 37°C till optical density reached 0.6. Protein expression was induced with 0.5 mM IPTG and incubated at 18°C overnight. Cells were harvested at 7,500 × g for 10 min at 4°C. Cell pellet was resuspended in PBS with 10% glycerol (volume equivalent to weight of cell pellet in grams) and either snap frozen in liquid nitrogen or used immediately.

Cells were resuspended in lysis buffer (50 mM sodium phosphate buffer, pH 7.4, 300 mM KCl, 5 % glycerol, 1 mM MgCl2, 10 mM B-ME, 0.1 mM ATP) supplemented with 30 mM Imidazole and protease inhibitor cocktail and lysed by 4-5 passages in an Emusiflux french press. Lysate was clarified at 1, 86, 000 × g for 1 hr at 4^°^C and passed through 0.45 μm membrane filter to further rid of particulate matter. His-trap column was equilibrated with 10 CV of lysis buffer, followed by lysate application at a flow rate of 1 mLmin^−1^. The column was washed with 10 CV of lysis buffer supplemented with 60 mM Imidazole and protein eluted in lysis buffer supplemented with 300 mM Imidazole. Protein was desalted, aliquoted, snap frozen and stored at — 80°C.

### Gliding assay

Glass coverslips (22 × 22 mm and 18 × 18 mm) were cleaned as follows: sonicated in 1:20 mucasol for 15 min, rinsed in distilled water for 2 min, sonicated in 100% ethanol for 10 min, rinsed in distilled water for 2 min, rinsed in double distilled water for 2 min and blow-dried with nitrogen. Flow channels were prepared by placing 1.5 mm parafilm strips on a cleaned 22 × 22 mm coverslip (about 3 mm apart 4 strips to prepare 3 channels) and covered with a 18 × 18 mm coverslip. This flow cell was placed on heat block maintained at 55° C to melt the parafilm thereby making channels water tight.

Aliquots of dynein complexes were subjected to microtubule sedimentation to get rid of dead/rigor binding motors before being used in gliding assays. Briefly, dynein complexes were incubated with unlabeled microtubules in dilution buffer (10 mM PIPES, pH 7.0, 50 mM potassium acetate, 4 mM MgSO4, 1 mM EGTA, 0.1 % Tween20, 10 μM taxol, 2 mM Mg-ATP, 10 mM dithiothreitol) for 10 min followed by centrifugation at 120,000 × g for 10 min at room temperature. Active dynein complexes retained in the supernatant were kept on ice until further use.

Flow channels were perfused with 2.5 mg mL^−1^ Protein A in double distilled water and incubated for 5 min followed by washing with dilution buffer. Solutions with varying concentrations (prepared in dilution buffer) of kinesin and dynein were incubated in these channels for 5 min and excess motors were washed out with dilution buffer. To block the rest of the surface, motor coated channels were incubated with 500 μg/mL casein in dilution buffer for 5 min followed by washing with dilution buffer. Double-stabilized microtubules prepared in motility buffer (dilution buffer supplemented with 40 mM glucose, 110 μgmL^−1^ glucose oxidase, 10μgmL^−1^ catalase) were incubated for 1 min and excess unbound microtubules were washed out with motility buffer.

Gliding microtubules were imaged on an inverted fluorescence microscope (Axiovert 200M, Zeiss) with a 40X, NA 1.3 oil immersion objective maintained at 27°C. Samples were illuminated with a metal arc lamp (Lumen 200, Prior Scientific) with an excitation 534/30emission 653/40 filter set in the optical path. 50-200 frames were acquired with an iXon Ultra EMCCD (Andor) camera with a 100 ms exposure time at a frame rate of 1 Hz. Images were acquired with MetaMorph (Universal Imaging). A minimum of three independent sets of gliding assays (per experimental condition) were performed to obtained statistically significant results.

### Data analysis

Microtubules were tracked with Fluorescence Image Evaluation Software for Tracking and Analysis (FIESTA) (Ruhnow et al., 2011) which automates Gaussian fitting of fluorescens signals to extract position coordinates. All tracks were manually curated to filter out tracks of microtubules with significant changes in length between successive frames as observed when two microtubules cross paths during gliding. Data analysis and visualization were carried out in MATLAB2014b (MathWorks, Natick, MA, USA). Instantaneous velocity was defined as the ratio of distance traversed by a microtubule and difference in time between consecutive frames. Velocities were smoothed with a rolling frame average over a window of five frames. Mean, median and quantiles of velocity distributions were calculated with in-built MATLAB functions.

Microtubule filament length was an important parameter that influenced gliding velocities (especially with dynein). Unless otherwise indicated, only 10 — 15 μm long microtubules (filtered from length values acquired from FIESTA) were used for analysis to obtain a more homogenous velocity distribution.

### Surface density estimation

Density of kinesin and dynein motors were estimated by taking into a consideration the protein concentration of motors perfused (7 μL) into a 2 × 18 × 3 mm^2^ flowcell: Highest dynein concentration of 55 μgmL^−1^ yielded a density of ~ 1280 μm^−2^. Motility measurements performed on axonemal dynein-coated surfaces have shown only 10% of the total motor population contribute to active motion (Kotani et al., 2007). Applying dynein concentrations of 55, 28, 11, 6, 3 and 1.5 μgml^−1^ were thus estimated to yield surface densities of 128, 64, 26, 13, 6 and 3 μm^−2^, respectively (Monzon and Scharrel et al., 2019). Similar assumptions were also made for estimated surface densities of kinesin.

The linear relationship between applied motor concentrations and surface density was confirmed by observing landing events of microtubules on motor coated surfaces (Katira et al., 2007; Agarwal et al., 2012). Number of microtubules landing on the surface were counted every 10 s for 100 frames. The plot of microtubule number vs time was fit to the curve *N*(*t*) = *N*_init_ + *N*_max_[1 – e^(*Rt*/*N*_max_)^], where N(t) is number of landed microtubules, *N*_init_ is number of microtubules non-specifically adsorbed to the surface, *N*_max_ is maximum number of microtubules in the field of view, *R* is the landing rate and *t* is time elapsed. Landing rate *R* derived from fit, microtubule area *A* (= Length of microtubule × 25 nm) and maximal diffusion-limited landing rate *Z* (assumed to be equal to landing rate of microtubules on very high densly coated motor surfaces) were plugged into the equation *σ* = – [ln(1 – *R/Z*)]/*A* to yield surface density *σ*.

Surface density of kinesin measured by the landing rate method was in agreement with the density estimated by the first method (7 μL of 6 μg mL^−1^ of kinesin solution yielded a density estimate of 102 μm^−2^ vs 100 μm^−2^ by the landing rate method).

### Simulation of kinesin and dynein models

Here we first describe details of the kinesin and dynein model and then present a simulation run. The dynein model was previously published in (Monzon and Scharrel et al., 2019) and the kinesin model in (Monzon and Scharrel et al., 2019; Klein et al., 2014). Here some parameter values were slightly changed and the models were slightly expanded. All expansions compared to the models published in (Monzon and Scharrel et al., 2019) are marked in the text. All parameter values are listed in table S1 including references to the literature whenever possible.

#### Gliding assay

The gliding assay is modeled as a one-dimensional system. This means, only one protofilament of the microtubule is taken into account. The microtubule itself is described as a rigid onedimensional object, which is situated at constant height above the one-dimensional surface (glass coverslip). Attached motors move the microtubule back and forth.

#### Attachment area

For the simulation, the motor density given by the experiment has to be converted to a number of motors able to attach to the microtubule. Having a microtubule with length *L*_MT_, we assume that all motors in the area *L*_MT_ × *L*_attach_ are able to attach to the microtubule. Thus the number of motors is:

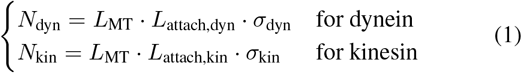

The widths of the attachment areas *L*_attach,dyn_ and *L*_attach,kin_ are the reach length of dynein and kinesin. In total *N*_kin_ + *N*_dyn_ motors are involved in the transport. In the simulation, *N*_kin_ + *N*_dyn_ motors are randomly distributed on the one-dimensional surface. To randomly distributed the motors, first the type of motor (kinesin or dynein) is thrown with probability *N*_kin_/(*N*_dyn_ + *N*_kin_) that the motor is a kinesin motor and with probability *N*_dyn_/(*N*_dyn_ + *N*_kin_) that it is a dynein motor. Next the position of the motor is randomly chosen taking hardcore interactions between the motors into account. If *D* = *L*_MT_/(*N*_dyn_ + *N*_kin_) is the mean distance between motors, 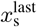 the position of the last set motor and *R_i_* the radii of the last, current and next set motor, the position of the currently set motor is uniformly distributed in the interval 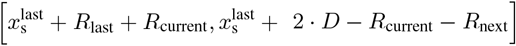.

#### Force calculation

Dynein and kinesin motors are modeled as linear springs. Therefore, the load force of a motor is calculated from the deflection of the motor. The deflection of a motor is the difference between the position of the head of the motor on the microtubule 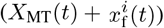^8^ and the position of the tail of the motor on the surface 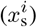. Thus the deflection is:

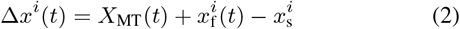

The load force of kinesin is divided into three regimes depending on its untensioned length *L*_0,kin_:

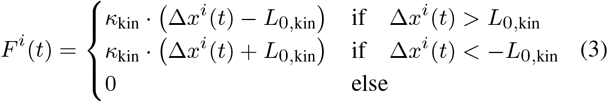

with *κ*_kin_ being the stiffness of kinesin. The load force of dynein is also divided into three regimes depending on *L*_0,dyn_ (= width of the deactivation region). However, unlike kinesin, dynein is not completely untensioned within *L*_0,dyn_. Within *L*_0,dyn_ the force is calculated using a smaller stiffness *κ*_1,dyn_:

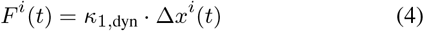

If dynein is stretched outside *L*_0,dyn_ the load force is

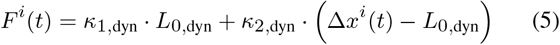

if Δ*x^i^*(*t*) > *L*_0,dyn_ and

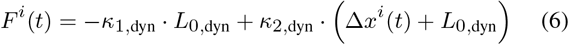

if Δ*x^i^*(*t*) < –*L*_0,dyn_. Thereby, is *κ*_2,dyn_ the dynein stiffness outside *L*_0,dyn_.

#### Attachment

If detached kinesin and dynein motors are under the microtubule (within the attachment area), they bind to the microtubule with the constant rates *k*_a,kin_ and *k*_a,dyn_, respectively.

#### Detachment

The detachment rates of attached kinesin and dynein motors depend on the load force of the motors. The detachment rate of kinesin increases exponentially with the load force and is symmetrically for forward (assisting force, *F^i^* ≤ 0) and backward load (resisting forces, *F^i^* > 0):

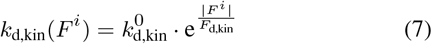

The detachment rate of dynein increases asymmetrically for forward (assisting force, *F^i^* > 0) and backward load (resisting forces, *F^i^* ≤ 0). It increases linearly with a steeper slope for forward load _1_ than for backward load:

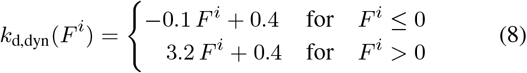

The dynein detachment behavior is taken from Cleary et al. (2014).

#### Stepping

Kinesin steps predominantly towards the microtubule plus end and active dynein towards the microtubule minus end. Kinesin steps in a force- and ATP-dependent manner, which is taken from Schnitzer et al. (2000). Under backward load forces smaller than the kinesin stall force (0 < *F^i^* < *F*_s,kin_) the stepping rate is:

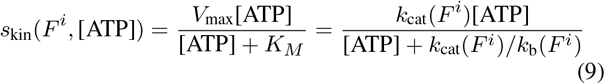

with *k*_cat_(*F^i^*) being the catalytic turnover rate constant and *k*_b_(*F^i^*) the second-order rate constant for ATP binding. Schnitzer et al. (2000) suggested a Boltzmann-type force dependence for the rate constants:

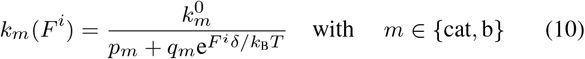

with 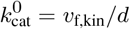. Thereby *v*_f,kin_ is the maximal kinesin stepping rate and *d* the step size. *q_m_, p_m_* with *q_m_* + *p_m_* = 1 and 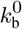 are taken from Schnitzer et al. (2000) and *δ* is determined by setting the stepping rate at stall force equal to 0.1 s^−1^:

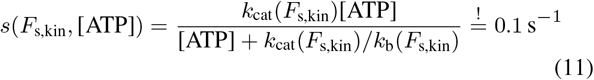

We solved this equation using MATLAB. For forward load forces (*F^i^* < 0) and the unloaded case (*F^i^* = 0), we apply *F^i^* =0 to Eqn. 9 and find the following Michaelis-Menten equation:

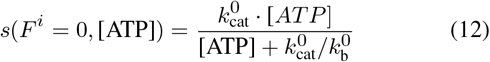

If kinesin is under very high backward load bigger than the kinesin stall force (*F^i^* > *F*_s,kin_), kinesin steps backwards (towards the microtubule minus end) with a low constant rate

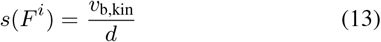

For <u>active dynein</u> the same force and ATP dependence of the stepping rate as for kinesin are applied with different parameter values. For backward forces below the stall force (0 > *F^i^* > –*F*_s,dyn_) Eqn. 9 and for forward load (*F^i^* > 0) Eqn. 12 are applied with 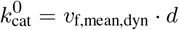 and *δ* determined by Eqn. 11 using *F*_s,dyn_. *q_m_, p_m_* and 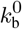 have the same values as for kinesin. Under backward load greater than the stall force (*F^i^* < – *F*_s,dyn_) active dynein steps backward with rate

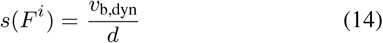

Another difference to kinesin is that for dynein the maximal velocity *v*_f,dyn_ is Gaussian distributed with the mean *v*_f,mean,dyn_, the standard deviation *σ_v_* and a cut at *v*_f,high_ and *v*_f,low_. Moreover, Mallik et al. (2004) found that dynein’s stall force is ATP dependent for low ATP concentrations. Here, we added the findings of Mallik et al. (2004) to the previously published dynein model (Monzon and Scharrel et al., 2019) and set the dynein stall force to

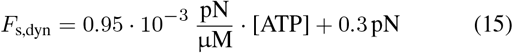

if [ATP] ≤ 1000 μM and to

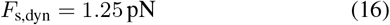

if [ATP] > 1000 μM.

Unlike the active dynein stepping predominately towards the microtubule minus end, the <u>passive dynein</u> diffuses in the harmonic potential of its linear spring. Thus the stepping rate is:

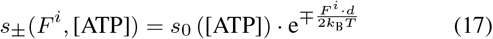

For the stepping rate of passive dynein in the unloaded case (*F^i^* = 0), we applied the same Michaelis-Menten equation as found for the unloaded case of directional stepping of active dynein (see Eqn. 12):

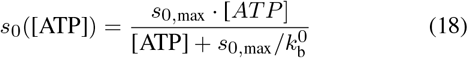

#### Mechanical dynein activation

When dynein attaches to the microtubule it is first in the passive state. When passive attached dynein is stretched more than *L*_0,dyn_, the deactivation region, it activates with rate *r*_a_ (Δ*x^i^*). The activation rate *r*_a_(Δ*x^i^*) depends on the deflection of the motor in an Arrhenius-like manner:

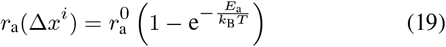

with *E*_a_ being the energy of the harmonic potential of the motor spring:

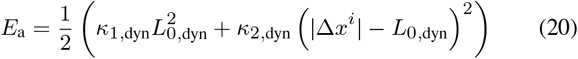

When active attached dynein is stretched less than *L*_0,dyn_, the deactivation region, it deactivates with the constant rate *r*_d_.

#### Roadblocks

For the simulations shown in Fig. 6 and S4, we added roadblocks to the previously published gliding assay model (Monzon and Scharrel et al., 2019). Here, we used rigor binding kinesin motor mutants (from now on referred to as roadblocks). The number of roadblocks on the microtubule is calculated as:

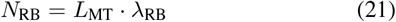

with λ_RB_ being the roadblock line density. The roadblocks are uniformly distributed on the one-dimensional microtubule (one protofilament), whereby hardcore interactions between the roadblocks were taken into account.

#### Gliding assay with seven protofilaments

To compare our results to cargo transport by multiple motors on several protofilaments (Ferro et al., 2019), we implemented a gliding assay with seven protofilaments in the presence of roadblocks. In this modification not only one protofilament is considered, like before, but seven protofilaments. The roadblocks and motors are randomly distributed on the seven protofilaments. If a motor encounters a roadblock or another motor on the same protofilament it cannot continue walking on this protofilament. However, when a motor encounters a roadblock or another motor protein which is on another protofilament, it is not influenced by this obstacle and continues walking normally. From previous single motor experiments, it is known that dynein frequently takes side steps (Reck-Peterson et al., 2006), while kinesin stays on the same protofilament (Ray et al., 1993). Here, we implemented a side-stepping rate *s*_side_ for dynein to change the protofilament. However, kinesin stays on the same protofilament as long as the kinesin motor is attached. Thus to circumvent a roadblock, dynein can use its side stepping ability while kinesin has to detach and reattach after the roadblock. The results of the gliding assay with seven protofilaments is shown in Fig. S3.

#### Simulation details

In the following, one run of the simulation is described.

##### Initialization

As a first step the one-dimensional surface is coated with motors. At the beginning the microtubule is situated at position *X*_MT_(*t* = 0) = 0 and no motor is attached to the microtubule.

##### Update

Using Gillespie’s first reaction sampling (Gillespie, 1977) the next event (attachment, detachment, stepping or (de)activation) is chosen. Then the chosen motor event is performed, i. e., the chosen motor is updated. After the motor event, the microtubule is moved to its nearest equilibrium position using a bisection search algorithm.

##### Measurement and output data

After the relaxation time trelax the microtubule position and instantaneous velocity are measured. Each second (like in the experiment) the time, the velocity and the position of the microtubule are measured and output as well as the number of attached motors (distinguishing between kinesin, active and passive dynein motors). To mimic the measurement uncertainty of the experiment, a white noise with standard derivation *σ*_pos_ is added to the actual simulation position.

The microtubule instantaneous velocity is measured from the last and current microtubule position (plus white noise) and the last and current time:

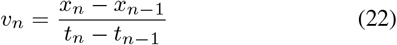

where *x*_*n*–1_ is the position at the last measurement time *t*_*n*–1_.

##### Termination

The complete simulation is terminated after *N*_samples_ program runs or if *N*_mes_ measurements were performed. A single program run is terminated either after a specific amount of measurements *n*_mes_ or after the simulation time *T*_end_, or when no motor is attached to the microtubule^9^. The reason for the last termination condition is that in the experiment a microtubule trajectory is also terminated once no motor is attached anymore. If no motor is attached anymore in the experiment, the microtubule diffuses vertically away from the surface and cannot be measured.

## Acknowledgements

We thank Andrew Carter and Max Schlager for kindly providing the plasmids encoding genes for H.sapiens dynein subunits and the multiBac plasmid;

## Contribution

Conceptualization: G.A.M., L. Scharrel, L. Santen, S.D.; Formal analysis: G.A.M., L. Scharrel, L. Santen, S.D.; Investigation: G.A.M., L. Scharrel, L. Santen, S.D.; Resources: L. Scharrel, S.D.; Data curation: G.A.M., L. Scharrel, L. Santen, S.D.; Writing - original draft: G.A.M.; Writing - review & editing: G.A.M., A.D., L. Santen, S.D.; Writing - experimental part of Material & Methods: A.D.; Visualization: G.A.M.; Supervision: L. Santen, S.D.; Project administration: L. Santen, S.D.; Funding acquisition: L. Santen, S.D.

## Supplementary

Insert the supplementary text here.

## Footnotes

1 The number of motors involved in the transport is determined by the motors being directly under the microtubule and in the immediate surrounding. Thus, first the density of the motors determines the actual number of motors involved in the transport and secondly the microtubule length. Here the microtubule length can be assumed to be constant since we always consider microtubules with approximately the same length, i.e. we only take into account microtubules with lengths in a certain interval. That is why the motor number and the motor density are used as synonyms.

2 We showed in our previous work (Monzon and Scharrel et al., 2019) that at low dynein-densities no directed dynein-driven transport occurs but the microtubule diffuses back and forth.

3 The harmonic potential comes from the motors springs.

4 The <u>mean</u> number of active attached dynein was greater than zero (0.43 active dynein motors). This means temporarily an active dynein is attached and drives the microtubule. That is why the median velocity of the dynein-driven state is not zero.

5 The reason is that motors step stochastically with different velocities. Which means there are not all aligned but their deflection is randomly distributed. See Material and Methods and previous literature for more details.

6 In the simulation instead of a roadblock concentration a roadblock line density is given.

7 and motors are not able to attach again

8 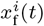 is the motor position on the microtubule which is zero when the motor is at the microtubule minus end and *L*_MT_ when the motor is at the microtubule plus end. *X*_MT_(*t*) is the position of the microtubule minus end in the overall coordination system of the surface. Therefore, the position of the head of the motor in the overall coordination system of the surface (coverslip) is 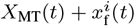.

9 At a time point greater than zero.

