## Supplementary Information for "Stable tug-of-war between kinesin-1 and cytoplasmic dynein upon different ATP and roadblock concentrations"

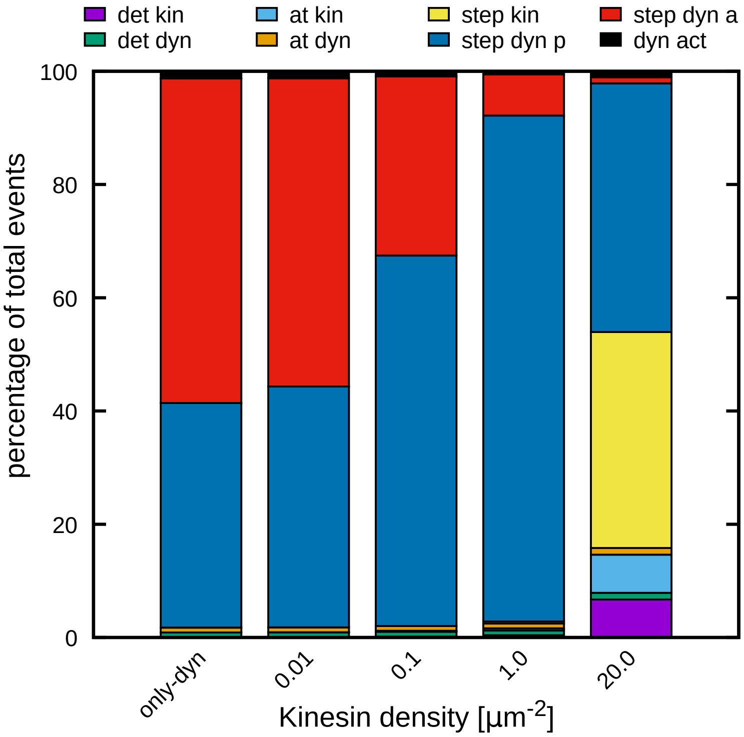


Figure S1: **In the balanced state mainly passive dynein stepping occurs.** We see for a constant dynein density of $\sigma_{\text{dyn}} = 64\,\mu\text{m}^{-2}$ the percentage of the occurring events as a function of the kinesin density. The percentage of the following events are presented: kinesin detachment (purple), dynein detachment (green), kinesin attachment (light blue), dynein attachment (orange), kinesin stepping (yellow), passive dynein stepping (blue), active dynein stepping (red) and dynein (de)activation (black). We see for the balanced state at $\sigma_{\text{kin}} = 1.0\,\mu\text{m}^{-2}$ that almost only passive dynein stepping occurrs. Kinesin almost does not step at all, and active dynein only steps little.

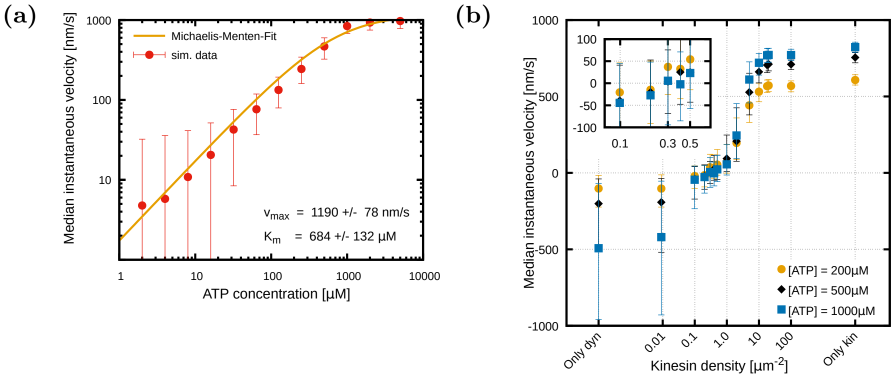


Figure S2: **Michaelis-Menten dependence for high dynein density and no significant shift of the balanced state for lower ATP concentrations.** a) Median instantaneous velocities with IQR as a function of ATP concentration for a unidirectional dynein assay simulation. Here a high dynein density of $\sigma_{\mathrm{dyn}} = 64\,\mu\mathrm{m}^{-2}$ was used and a microtubule length of $L_{\mathrm{MT}} = 25\,\mu\mathrm{m}$. For a high dynein density the simulation data could be fit with a Michaelis-Menten equation $V_{\max} \times [\mathrm{ATP}]/(K_m + [\mathrm{ATP}])$ with $V_{\max} = 1190 \pm 78\,\mathrm{nm/s}$ and $K_m = 684 \pm 132\,\mu\mathrm{M}$. b) Median instantaneous velocities with IQR for bidirectional gliding assays at different ATP concentration. As in Fig. 5C of the main text a constant intermediate dynein density of $\sigma_{\mathrm{dyn}} = 18\,\mu\mathrm{m}^{-2}$ was used for all ATP concentrations. We see that for lower ATP concentrations the balanced state slightly shifts towards lower kinesin densities. However, the shift is in the order of magnitude of the fluctuations arising from a fluctuating number of motors involved in the transport process of microtubule gliding assays. Thus, no significant dependence on the ATP concentration could be observed.

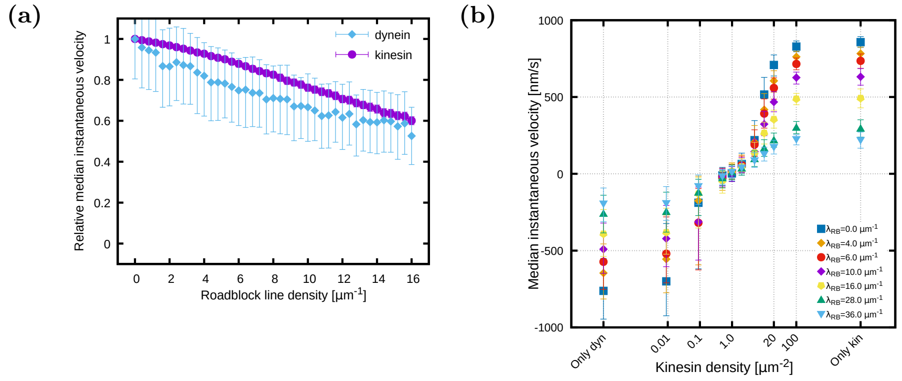


Figure S3: **In several protofilament simulations dynein is as affected by roadblocks as kinesin, however the balanced state stays at the same kinesin density for all raodblock densities.** a) Relative median instantaneous velocities as a function of the roadblock line density for unidirectional dynein (light blue) and kinesin (purple) assay simulations. Here, a modified simulation was used. In this simulation instead of one protofilament, seven protofilaments were modeled. Additionally, to change a protofilament, for dynein a side-stepping rate of $s_{\text{side}} = 4\,\text{s}^{-1}$ was implemented. Unlike dynein, kinesin is not able to side step and change the protofilament. Like in the simulation shown in Fig. 6a of the main text, the microtubule length was $L_{\text{MT}} = 25\,\mu\text{m}$ and the kinesin and dynein densities were $\sigma_{\text{kin}} = \sigma_{\text{dyn}} = 50\,\mu\text{m}^{-2}$. We see that in this case both motors are less affected by roadblocks than in the one-protofilament simulation (see Fig. 6a of the main text). Moreover, dynein is as affected by roadblocks as kinesin. b) Median instantaneous gliding assay velocities as a function of varying kinesin density for bidirectional gliding simulations at different roadblock line densities. As in a) the modified simulation with seven protofilaments was used. Like in the simulation shown in Fig. 6c of the main text, the microtubule length was $L_{\text{MT}} = 25\,\mu\text{m}$ and the constant dynein density was $\sigma_{\text{dyn}} = 50\,\mu\text{m}^{-2}$. We see that for all roadblock line densities the balanced state stays at the same kinesin density.

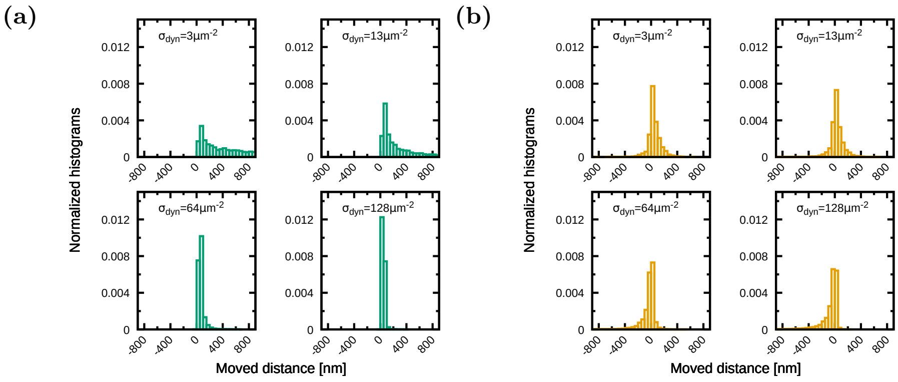


Figure S4: **Attached kinesin and dynein motors in the balanced state move very little.** Histograms of the moved distance of a) kinesin and b) dynein. The used dynein densities are depicted in the subfigures and the kinesin density was $\sigma_{\mathrm{kin}} = 1.0\,\mu\mathrm{m}^{-2}$ for all shown histograms. The kinesin density of $\sigma_{\mathrm{kin}} = 1.0\,\mu\mathrm{m}^{-2}$ is the balanced state kinesin density for a dynein density of $\sigma_{\mathrm{dyn}} = 64\,\mu\mathrm{m}^{-2}$. For the balanced state ($\sigma_{\mathrm{dyn}} = 64\,\mu\mathrm{m}^{-2}$) we see that the moved distance of kinesin a) and dynein b) is peaked around zero. This means in the balanced state, motors are strongly localized.

| Description: | Value | Reference: |
|---|---|---|
| **Common parameters:** | | |
| Microtubule length | $L_{\mathrm{MT}} = 25\,\mu\mathrm{m}$, if not further specified | In the range of the experiment |
| Dynein surface density | $\sigma_{\mathrm{dyn}}$, varied in the experiments and simulations | Estimated from the experimental dynein solution. |
| Kinesin surface density | $\sigma_{\mathrm{kin}}$, varied in the experiments and simulations | Estimated from the experimental kinesin solution. |
| Stepsize of kinesin and dynein | $d = 8\,\mathrm{nm}$ | [4, 23, 13] |
| ATP concentration | $[\mathrm{ATP}] = 2000\,\mu\mathrm{M}$, if not further specified | Same value as in the experiment. |
| Temperature | $T = 300\,\mathrm{K}$ | Like in the experiment |
| Unloaded second-order rate constant for ATP binding | $k_{\mathrm{b}}^{0} = 1.3\,\mu\mathrm{M}^{-1}\mathrm{s}^{-1}$ | [19] |
| Fraction of unloaded catalytic cycle | $q_{\mathrm{cat}} = 6.2$, $p_{\mathrm{cat}} = 1 - q_{\mathrm{cat}}$ | [19] |
| Fraction of unloaded catalytic cycle | $q_{\mathrm{b}} = 4.0$, $p_{\mathrm{b}} = 1 - q_{\mathrm{b}}$ | [19] |
| **Dynein:** | | |
| Motor radius | $R^{i}_{\mathrm{dyn}} = 24\,\mathrm{nm}$ | Approximated from EM images of [22, 18] |
| Length of deactivation area | $L_{0,\mathrm{dyn}} = 30\,\mathrm{nm}$ | Unknown |
| Width of attachment area | $L_{\mathrm{attach,dyn}} = 30\,\mathrm{nm}$ | Dynein length: Approximated from EM images of [22, 18] |
| Stiffness within $L_0$ | $\kappa_{1,\mathrm{dyn}} = 0.019272\,\mathrm{pN/nm}$ | Unknown. Adjust in order to obtain the same number of steps until reaching stall as with the higher $k_{2,\mathrm{dyn}}$ used in our previous paper (Monzon and Scharrel et. al. 2019) |
| Stiffness beyond $L_0$ | $\kappa_{2,\mathrm{dyn}} = 0.065\,\mathrm{pN/nm}$ | Lower than in our previous publication (Monzon and Scharrel et. al. 2019). Here adjusted to the value of Ohashi et. al. 2019 |

| | | |
|---|---|---|
| Stall force | $F_{\mathrm{s,dyn}} = 1.25\,\mathrm{pN}$ | Same as [16, 12]. Same order of magnitude as [13, 1, 7, 17] |
| Attachment rate | $k_{\mathrm{a,dyn}} = 0.2\,\mathrm{s}^{-1}$ | Unknown |
| Mean forward velocity | $v_{\mathrm{f,mean,dyn}} = 1300\,\mathrm{nm/s}$ | [22] (Motility assay) |
| Backward velocity | $v_{\mathrm{b,dyn}} = 15\,\mathrm{nm/s}$ | [6] (yeast dynein) |
| Standard deviation of max. velocity distribution | $\sigma_v = 1500\,\mathrm{nm/s}$ | Relatively wide velocity distributions were measured in [14, 18], too. |
| Left velocity border | $v_{\mathrm{f,low}} = 300\,\mathrm{nm/s}$ | |
| Right velocity border | $v_{\mathrm{f,high}} = 2300\,\mathrm{nm/s}$ | |
| Diffusion rate | $s_{0,\mathrm{max}} = 92.9\,\mathrm{s}^{-1}$, $s_0([\mathrm{ATP}] = 2000\,\mu\mathrm{M}) = 85\,\mathrm{s}^{-1}$ | From [15] (Fit of experimental data) |
| Activation rate constant | $r_a^0 = 40\,\mathrm{s}^{-1}$ | Unknown |
| Deactivation rate | $r_d = 1\,\mathrm{s}^{-1}$ | Unknown |
| Motor radius | $R^i_{\mathrm{dyn}} = 24\,\mathrm{nm}$ | Approximated from EM images of [22, 18] |
| **Kinesin:** | | |
| Motor radius | $R_{\mathrm{kin}} = 4\,\mathrm{nm}$ | Same order of magnitude as [21] |
| Untensioned length | $L_{0,\mathrm{kin}} = 30\,\mathrm{nm}$ | [11] |
| Width of attachment area ≈ twice kinesin contour length | $L_{\mathrm{attach,kin}} = 100\,\mathrm{nm}$ | Same order of magnitude as [11, 8, 20] |
| Stiffness | $\kappa_{2,\mathrm{kin}} = 3.0 \cdot 10^{-4}\,\mathrm{kg/s}^2$ | [10, 9, 16] |
| Stall force | $F_{\mathrm{s,kin}} = 6\,\mathrm{pN}$ | [23, 12] |
| Detachment force | $F_{\mathrm{d,kin}} = 6\,\mathrm{pN}$ | [12] |
| Force free detachment rate | $k^0_{\mathrm{d,kin}} = 0.66\,\mathrm{s}^{-1}$ | Taken from [16]. Same order of magnitude as [19] |
| Attachment rate | $k_{\mathrm{a,kin}} = 20\,\mathrm{s}^{-1}$ | |
| Forward velocity | $v_{\mathrm{f,kin}} = 1000\,\mathrm{nm/s}$ | [4, 23] |
| Backward velocity | $v_{\mathrm{b,kin}} = 6\,\mathrm{nm/s}$ | Same order of magnitude as [3] |
| **Simulation parameters:** | | |
| Relaxation time (=time until measurements start) | $t_{\mathrm{relax}} = 20\,\mathrm{s}$ | |
| Total simulation time | $t_{\mathrm{end}} = 220\,\mathrm{s}$ | |
| Number of runs | $N_{\mathrm{samples}} = 15$ (Fig. 4) $N_{\mathrm{samples}} = 20$ (Fig. 5&6) | |

| | | |
|---|---|---|
| Number of measurements in one run | $n_{\mathrm{mes}} = 200$ | |
| Total number of measurements of all runs together | $N_{\mathrm{mes}} = 3000$ (Fig. 4)<br>$N_{\mathrm{mes}} = 4000$ (Fig. 5&6) | |
| Standard deviation of MT position | $\sigma_{\mathrm{Pos}} = 30\,\mathrm{nm}$ | measurement uncertainty from experiment |
| **Roadblocks:** | | |
| Roadblocks line density | $\lambda_{\mathrm{RB}} = 0 - 16\,\mu\mathrm{m}^{-1}$ (stated in the figures) | Range chosen similar to [5] |
| Number of Roadblocks on the microtubule | $N_{\mathrm{RB}}$ | Calculated from the roadblock line density $\lambda_{\mathrm{RB}}$ and the microtubule length $L_{\mathrm{MT}}$ |
| Radius of roadblocks (rigor binding kinesin motor mutants) | $R_{\mathrm{RB}} = 4\,\mathrm{nm}$ | As kinesin motor |
| **Modified simulation with several protofilaments:** | | |
| Number of protofilaments | $N_{\mathrm{L}} = 7$ | Optimized until obtaining the kinesin result shown by [5] for multiple kinesin motors |
| Dynein side stepping rate | $s_{\mathrm{side}} = 4\,\mathrm{s}^{-1}$ | Same order of magnitude as calculated from results of [5] (mammalian) and [2] (yeast) |

Table S1: Simulation parameter at the default ATP concentration of $[\mathrm{ATP}] = 2000\,\mu\mathrm{M}$.
